# H3.1K27M-induced misregulation of the TSK/TONSL-H3.1 pathway causes genomic instability

**DOI:** 10.1101/2024.12.09.627617

**Authors:** Wenxin Yuan, Yi-Chun Huang, Chantal LeBlanc, Axel Poulet, Devisree Valsakumar, Josien C. van Wolfswinkel, Philipp Voigt, Yannick Jacob

**Author notes:** These authors contributed equally to this manuscript (Wenxin Yuan, Yi-Chun Huang).

## Abstract

The oncomutation lysine 27-to-methionine in histone H3 (H3K27M) is frequently identified in tumors of patients with diffuse midline glioma-H3K27 altered (DMG-H3K27a). H3K27M inhibits the deposition of the histone mark H3K27me3, which affects the maintenance of transcriptional programs and cell identity. Cells expressing H3K27M are also characterized by defects in genome integrity, but the mechanisms linking expression of the oncohistone to DNA damage remain mostly unknown. In this study, we demonstrate that expression of H3.1K27M in the model plant *Arabidopsis thaliana* interferes with post-replicative chromatin maturation mediated by the H3.1K27 methyltransferases ATXR5 and ATXR6. As a result, H3.1 variants on nascent chromatin remain unmethylated at K27 (H3.1K27me0), leading to ectopic activity of TONSOKU (TSK), which induces DNA damage and genomic alterations. Elimination of TSK activity suppresses the genome stability defects associated with H3.1K27M expression, while inactivation of specific DNA repair pathways prevents survival of H3.1K27M-expressing plants. Overall, our results suggest that H3.1K27M disrupts the chromatin-based mechanisms regulating TSK/TONSL activity, which causes genomic instability and may contribute to the etiology of DMG-H3K27a.

## Introduction

DMG-H3K27a, a type of brain cancer that mostly affects children, is characterized by a very poor prognosis with fewer than 10% of patients surviving more than two years after diagnosis (*1*). Approximately 80% of DMG patients are carriers of a somatic K27M mutation in one of the histone H3 genes (*2, 3*). The H3K27M mutation can occur in genes encoding different histone H3 variants: replication-dependent H3.1 or H3.2 (H3.1/H3.2 variants hereafter referred to as H3.1), and replication-independent H3.3 (*3, 4*). Initial work has shown that expression of H3.1K27M or H3.3K27M leads to a decrease of histone H3 lysine 27 tri-methylation (H3K27me3) by inhibiting the activity of the H3K27 methyltransferase POLYCOMB REPRESSIVE COMPLEX 2 (PRC2) (*5–8*). In cancer cells, H3.1K27M and H3.3K27M are both expressed amid a much larger contingent of wild-type H3.1 and H3.3 proteins, but the mutated histones inhibit PRC2 in a dominant-negative manner (*5, 7*), which accounts for the drastic loss of H3K27me3 observed in DMG-H3K27a cells.

A large majority of the work on oncogenic H3K27M mutations has centered on the consequences of decreased H3K27me3 levels and the subsequent effects on transcriptional regulation. However, the disruption of other cellular activities by H3K27M may have been overlooked, as PRC2 activity in mammals and *Drosophila melanogaster* is responsible for all levels of methylation at H3K27 (*9–11*). Mono- and di-methylation at H3K27 (H3K27me1/2), which are together much more abundant than H3K27me3 in mouse embryonic stem cells (*10*), have cellular functions that are independent of H3K27me3 (*10, 12, 13*). In line with this, it has been confirmed that levels of H3K27me1/2 at specific loci are also reduced by the H3K27M oncomutation in H3K27M-expressing cell lines (*14*). Consequently, there is a major increase in unmethylated histone H3 at K27 (H3K27me0), to the point where it can become the most dominant form of H3K27 (*15*). Importantly, PRC2 inhibition by H3K27M leads to a large increase of K27me0 on both H3.1 and H3.3 variants (*15*).

Lysine 27 of newly synthesized H3.1 proteins is unmethylated prior to incorporation of H3.1 into chromatin during DNA replication (*16, 17*). Recently, it was shown in the model plant *Arabidopsis thaliana* (Arabidopsis) that H3.1K27me0 is specifically required for the recruitment of the conserved DNA repair protein TONSOKU (TSK; known as TONSL in animals) to replication forks (*18*). In mammals, TONSL has been shown to initiates homologous recombination-mediated resolution of stalled or broken replication forks (*19–22*). DNA repair via TSK/TONSL needs to be tightly regulated in dividing cells of plants and animals by histone methyltransferases, which methylate post-replicative chromatin to prevent TSK/TONSL activity outside of stalled or broken replication forks (*18, 23*). In the absence of the post-replicative chromatin maturation step that inhibits TSK/TONSL, genome instability is observed in a TSK/TONSL-dependent manner (*18*). Chromatin-based regulation of TSK in plants has been shown to depend on the redundant activity of the H3.1K27 mono-methyltransferases ATXR5 and ATXR6 (ATXR5/6) (*24, 25*), which mono-methylate H3.1K27 to prevent binding of TSK to H3.1 through its conserved tetratricopeptide repeat (TPR) domain (*18, 25*). In Arabidopsis, where unregulated TSK activity has been extensively studied, this results in many phenotypes, including widespread genomic amplification of heterochromatic sequences (hereafter referred to as heterochromatin amplification), DNA damage, and loss of transcriptional silencing (*18, 25–27*).

Genomic instability is a defining characteristic of many cancer cells (*28*), including H3.1K27M and H3.3K27M tumors (*29–33*). The frequent co-occurrence of specific secondary mutations associated with H3.1K27M and H3.3K27M (*32*), and the defects in genome integrity in cancer cells harboring these mutations, suggest a model where expression of H3K27M generates a DNA error-prone environment that can induce and/or accelerate tumorigenesis (*33*). Accordingly, specific DNA repair pathways and DNA repair pathway choice have been shown to be affected by H3K27M (*30, 33*), but a direct link between the oncohistone and misregulation of DNA repair proteins has not yet been established. In this work, we demonstrate that expression of H3.1K27M disrupts the regulation of the TSK/TONSL-H3.1 DNA repair pathway. Using Arabidopsis as a model system for K-to-M mutations on H3 (*34*), we show that expression of H3.1K27M induces DNA damage. This effect of H3.1K27M is due to its ability to block the activity of ATXR5/6, which results in an increase of H3.1K27me0 in chromatin. Similarly to *atxr5/6* mutants, increased levels of H3.1K27me0 induce heterochromatin amplification and loss of transposon silencing. By inactivating TSK in H3.1K27M-expressing plants, we were able to confirm that ectopic activity of the TSK-H3.1 DNA repair pathway is responsible for disrupting genome integrity. In addition, loss of the DNA repair proteins MRE11 and RAD51 in H3.1K27M-expressing plants results in synthetic lethality, thus suggesting specific vulnerabilities in cells expressing H3.1K27M. We discuss the implications of these results obtained in the Arabidopsis model and how they could translate to a better understanding of the etiology of cancers characterized by H3.1K27M expression.

## Results

### H3.1K27M expression induces genomic instability in Arabidopsis

To explore the biological mechanisms affected by an H3K27M mutation, we created transgenic Arabidopsis plants expressing H3.1K27M under a native H3.1 promoter (H3.1_prom_::H3.1K27M) in the wild-type Columbia-0 (Col-0) background, which also expresses a normal set of endogenous histone H3.1 from five H3.1 genes (*35*). As a control, we also created transgenic plants expressing wild-type H3.1 (H3.1_prom_::H3.1 WT). RNA-seq analysis showed that H3.1K27M transcripts corresponded to ∼20% of all H3.1 transcripts (Fig. 1A), indicating that a majority of H3.1 proteins in the transgenic plants are wild type. This approach provides the opportunity to evaluate the phenotypic and molecular effects caused by H3.1K27M expression in a model system where H3-variant-specific effects on genome stability and DNA repair have been previously characterized (*18*).

**Figure 1.**
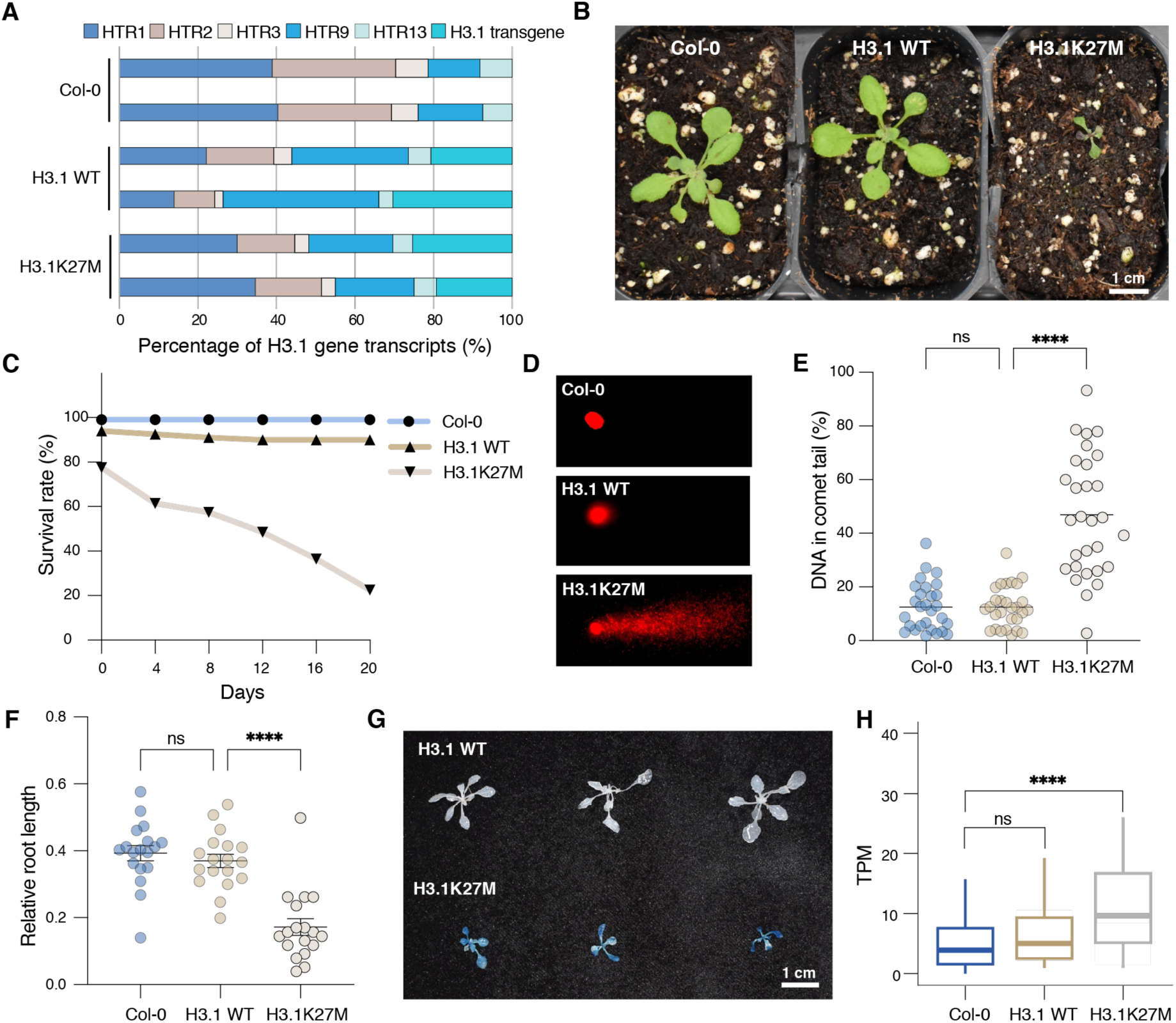
H3.1K27M expression induces developmental defects and DNA damage in Arabidopsis. **A** Relative abundance of H3.1 gene transcripts. **B** Morphological phenotypes of Col-0, H3.1 WT and H3.1K27M plants. **C** Survival rates of Col-0, H3.1 WT and H3.1K27M plants. **D** Representative comet assay images. **E** Quantification of DNA percentage in the comet tails. Horizontal bars indicate the mean. Welch’s ANOVA followed by the Dunnett’s T3 test: **** *p* < 0.0001, ns = not significantly different. **F** Relative root length of seedlings grown on ½ MS plates with 100 µg/ml MMS compared to the average root length of seedlings grown on plates without MMS. Each dot represents one individual T1 plant. Horizontal bars indicate the mean. SEM is shown. One-way ANOVA with Tukey’s multiple comparison test: **** *p* < 0.0001, ns = not significantly different. **G** Representative image of histochemical staining of reporter plants for GUS activity. Blue areas indicate that a functional reporter gene has been restored via a somatic recombination event. **H** RNA-seq data showing relative transcript levels of 158 DNA damage response genes (Table S1) measured by transcripts per million (TPM). Wilcoxon test with Bonferroni correction: **** *p* < 0.0001, ns = not significantly different.

The H3.1K27M transgenic plants demonstrated significant developmental defects compared to Col-0 and wild-type H3.1 controls (H3.1 WT), exhibiting stunted growth and much smaller and narrower leaves compared to the control plants (Fig. 1B and fig. S1A). The germination and survival rates of H3.1K27M plants were severely affected, with most plants dying before day 20 (Fig. 1C). In addition, the H3.1K27M plants displayed an increase in anthocyanin content, which is commonly associated with cellular stress (fig. S1B) (*36*). To assess for the presence of DNA damage, we conducted comet assays and found a significant increase in tail DNA percentage for H3.1K27M-expressing plants compared to Col-0 and H3.1 WT controls, which is indicative of DNA breaks (Fig. 1, D and E). We also found that H3.1K27M plants were hypersensitive to the genotoxic effects of methyl methanesulfonate (MMS) (Fig. 1F and fig. S1C). We then used the disrupted beta-glucuronidase (*uidA*/*GUS*) system to directly assess the frequency of homologous recombination events in our transgenic plants (*37*). In this system, homology-directed repair is required to reconstitute a gene that can produce a functional GUS protein. Our results showed that GUS activity was much stronger in H3.1K27M transgenic plants, indicating higher levels of homology-directed repair at the *GUS* gene (Fig. 1G). In accord with these results, RNA-seq experiments in Col, H3.1 WT and H3.1K27M plants showed that many DNA damage response genes are upregulated when H3.1K27M is expressed (Fig. 1H and Table S1). Taken together, these results indicate that expression of H3.1K27M in Arabidopsis generates DNA damage.

### H3.1K27M expression inhibits the histone methyltransferase activity of PRC2 and ATXR5/6

In human cells, H3K27M mutations interfere with PRC2 in a dominant-negative manner, which prevents the spreading of H3K27me3 (*5, 7*). Given the high degree of conservation between plant and animal histone H3 variants and PRC2 complexes (*35, 38*), we hypothesized that expression of H3K27M in plants would also block PRC2 activity. To verify this, we performed immunostaining and observed a drastic decrease of H3K27me3 in nuclei from H3.1K27M-expressing plants compared to Col-0 and H3.1 WT (Fig. 2A). We also assessed H3K27me3 levels by chromatin immunoprecipitation sequencing combined with exogenous chromatin spike-in (ChIP-Rx) to normalize the H3K27me3 signal between samples. Profiles for H3K27me3 over protein-coding genes showed a large decrease in H3K27me3 in H3.1K27M plants compared to Col-0 and H3.1 WT in both high-and low-expressing genes (Fig. 2B and fig. S2A). Together, these results suggest that expression of the oncohistone H3.1K27M disrupts PRC2 activity in plants similarly to human cells.

**Figure 2.**
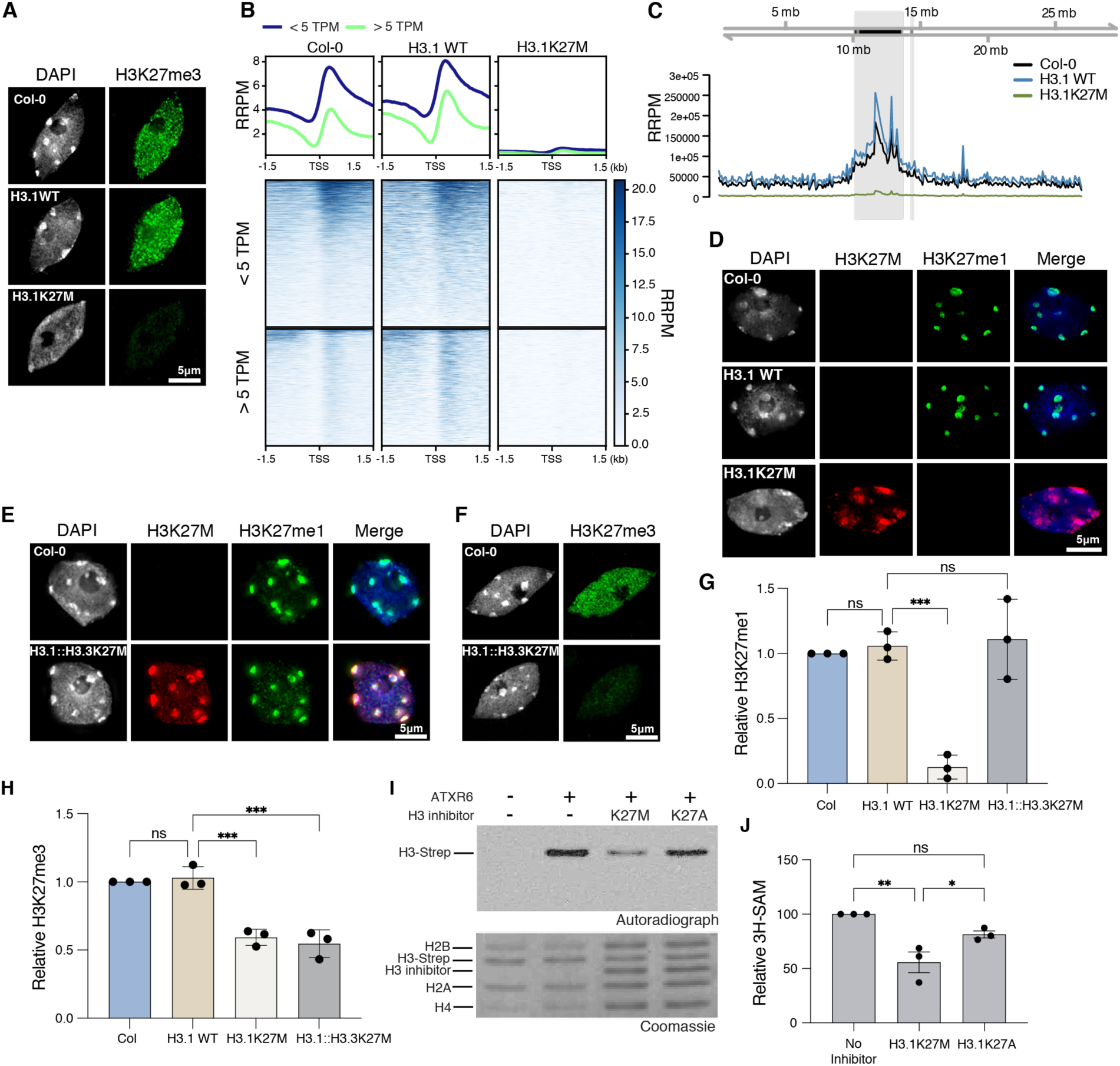
H3.1K27M-expressing plants exhibit loss of ATXR5/6-mediated H3K27me1. **A** Staining of leaf nuclei with an anti-H3K27me3 antibody and DAPI. **B** ChIP-seq profiles and heatmaps of normalized H3K27me3 signal over protein-coding genes grouped by level of expression (< 5 and > 5 TPM) in reference-adjusted reads per million (RRPM). TSS, transcription start site. **C** ChIP-Rx normalized H3K27me1 signal using 100 kb windows over chr 5. The pericentromeric region is shown in gray. **D** to **F** Staining of leaf nuclei with anti-H3K27M and anti-H3K27me1 (D and E) or anti-H3K27me3 antibodies (F) and DAPI. **G** and **H** Relative Western blot quantification of H3K27me1 (G) and H3K27me3 (H) levels in leaf total histones. One-way ANOVA with Tukey’s multiple comparison test: *** *p* < 0.001, ns = not significantly different. **I** Representative *in vitro* histone methyltranferase assay using ATXR6 and recombinant nucleosomes containing plant H3.1-strep. Nucleosomes containing H3K27M or H3K27A (H3 inhibitor) were added to the reactions. **J** Quantification of the relative amount of ^3^H-SAM incorporated into the H3.1-strep substrate. Each dot represents one independent experiment. Bars represent the mean. SEM is shown. One-way ANOVA with Tukey’s multiple comparison test: ** *p* = 0.0040, * *p* = 0.0459, ns = not significantly different.

In plants, H3K27 methylation is also catalyzed by the SET domain-proteins ATXR5/6, which specifically mono-methylate the H3.1 variant (*24, 25*). K-to-M mutations on histone H3 has been shown to inhibit the activity of multiple histone methyltransferases (*39*), so we predicted that expression of H3.1K27M in Arabidopsis may not only inhibit PRC2, but also ATXR5/6. To validate this hypothesis, we first performed ChIP-Rx analysis and observed a large decrease in H3K27me1 signal over heterochromatin in H3.1K27M-expressing plants compared to Col-0 and H3.1 WT plants (Fig. 2C and fig. S2B). Immunofluorescence staining using an H3K27M antibody showed that the mutated histone is enriched at chromocenters (Fig. 2D), as expected, because H3.1 proteins are concentrated within heterochromatic domains in somatic cells of Arabidopsis (*40, 41*). Staining with an H3K27me1 antibody showed the characteristic enrichment of this histone mark at chromocenters in Col-0 and H3.1 WT plants (*42*), but a drastic reduction in H3.1K27M-expressing plants (Fig. 2D). To demonstrate that the loss of H3K27me1 was due to ATXR5/6 inhibition, we generated transgenic plants expressing H3.1_prom_::H3.3K27M in the wild-type Col-0 background. As ATXR5/6 are unable to interact with H3.3 (*24*), we hypothesized that they would not be inhibited by H3.3K27M, but that PRC2 complexes would be negatively impacted as they can interact with both H3.1 and H3.3 in plants. As expected, we found that the level of H3K27me1 in H3.1_prom_::H3.3K27M plants was similar to Col-0, indicating that the presence of H3.3K27M did not inhibit ATXR5/6 (Fig. 2E). We did, however, find that the level of H3K27me3 was reduced in these plants (Fig. 2F), thus confirming that PRC2 activity in Arabidopsis is disrupted by H3.3K27M. Consistent with the immunostaining experiments, we observed a large decrease of H3K27me1 in H3.1K27M-expressing plants, but not in H3.1_prom_::H3.3K27M plants, in total histone samples extracted from leaves (Fig. 2G and fig. S2C). In contrast, a reduction in H3K27me3 was seen in both H3.1K27M and H3.1_prom_::H3.3K27M plants (Fig. 2H and fig. S2C).

Previous studies have shown that H3K27M acts in a dominant-negative manner by directly inhibiting the catalytic subunit (i.e., EZH2) of PRC2 (*5, 7, 43, 44*). Based on this, we hypothesized that the effect of H3.1K27M on H3K27me1 levels may work through a similar direct inhibitory mechanism on ATXR5/6. To test this, we performed *in vitro* histone methyltransferase (HMT) assays with and without H3.1K27M nucleosomes serving as potential ATXR6 inhibitors. Our results showed decreased levels of methylated H3.1 substrates when H3.1K27M nucleosomes were present in the reaction (Fig. 2, I and J). In contrast, H3.1K27A (lysine 27-to-alanine) had a smaller inhibitory effect than H3.1K27M on ATXR6 (Fig. 2, I and J), which is comparable to the effects of these two H3.1 mutants on mammalian PRC2 activity *in vitro* (*45*). Overall, our data indicate that, in the presence of H3.1K27M, ATXR5/6-mediated mono-methylation of H3.1K27 is inhibited, leading to widespread loss of H3.1K27me1 in the Arabidopsis genome.

### H3.1K27M induces genomic instability defects similarly to *atxr5/6* mutants

Loss of H3.1K27me1 in *atxr5/6* mutants leads to various heterochromatic phenotypes, including chromocenter decondensation, heterochromatin amplification (i.e., an increase in the copy number of transposons and other repetitive elements) and loss of transposon silencing (*25–27*). As H3.1K27M-expressing plants show a massive reduction of H3K27me1 (Fig. 2, C and D), we hypothesized that they would have heterochromatic defects similar to *atxr5/6* mutants. DAPI staining of nuclei from H3.1K27M-expressing plants revealed a hollowed sphere conformation of chromocenters characteristic of nuclei from *atxr5/6* mutant plants (Fig. 3, A and B) (*46*). Flow cytometry analyses showed that H3.1K27M plants exhibit broader peaks corresponding to endoreduplicated nuclei, a phenotype associated with heterochromatin amplification and also observed in *atxr5/6* mutants (Fig. 3, C and D) (*26*). In accord with this, whole-genome sequencing of H3.1K27M-expressing plants revealed that heterochromatic regions are amplified similarly to *atxr5/6* mutants (Fig. 3E). Finally, RNA-seq analysis indicated transcriptional reactivation of transposable elements (TEs) normally silenced in a Col-0 background, like in *atxr5/6* mutants (Fig. 3F).

**Figure 3.**
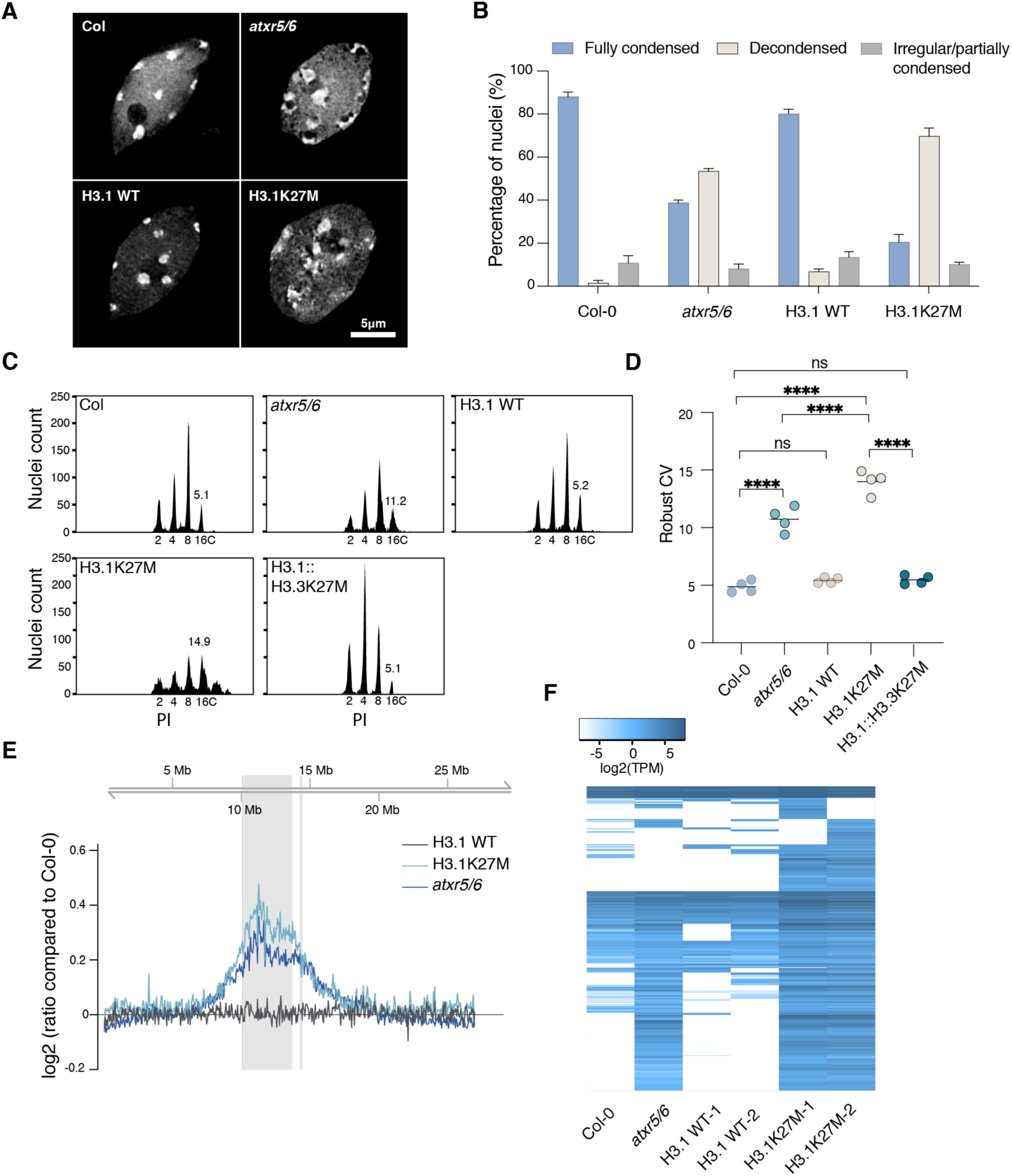
H3.1K27M induces heterochromatin defects similarly to *atxr5/6* mutants. **A** Leaf interphase nuclei stained with DAPI. **B** Quantification of chromocenter appearance of nuclei from experiment in panel A. **C** Flow cytometry profiles of leaf nuclei stained with propidium iodide (PI). Ploidy levels of the nuclei are shown below the peaks. The numbers above the 16C peaks indicate the robust coefficient of variation (CV). **D** Robust CV quantification of 16C peaks. Each dot represents an individual plant. The mean is shown. One-way ANOVA with Tukey’s multiple comparison test: **** *p* < 0.0001, ns = not significantly different. **E** Chromosomal view (Chr 5) of DNA-seq reads. The pericentromeric region is highlighted in gray. **F** Heat map showing the relative expression levels of TEs induced in H3.1K27M plants (Table S2).

In all our assays (Fig. 3, A to F), we observed an increase in the magnitude of heterochromatic defects in H3.1K27M plants compared to *atxr5/6* mutants. This observation is in line with the morphological comparison of these two genetic backgrounds, with *atxr5/6* mutants showing only a mild leaf developmental phenotype while H3.1K27M-expressing plants are drastically smaller and struggle to make it to the adult stage (fig. S3A). The well-characterized *atxr5/6* double mutant used in our experiments is a hypomorphic mutant, as *ATXR6* expression from the mutant allele is only reduced compared to wild-type plants (*25*). We use this particular mutant for our studies because complete elimination of ATXR5 and ATXR6 activity is lethal in Arabidopsis (*47*). The phenotypic variation between H3.1K27M-expressing plants and *atxr5/6* mutants may be linked to the difference in H3.1K27me1 levels between the two genetic backgrounds. This hypothesis is supported by Western blot analyses showing that H3.1K27me1 levels are much lower in H3.1K27M-expressing plants than *atxr5/6* mutants in two-week old plants (fig. S3B and C). In summary, our results indicate that expression of H3.1K27M mimics the phenotypes of *atxr5/6* mutants, and that *in vivo* levels of H3.1K27me1 may dictate the severity of these phenotypes.

### H3.1K27M induces genomic instability in a TSK-dependent manner

In plants, ATXR5/6 methylate newly synthesized H3.1 variants inserted on chromatin to induce post-replicative chromatin maturation, a key regulatory step that prevents the activity of the DNA repair protein TSK by inhibiting its binding to chromatin (*18, 24, 25, 48*). TSK is recruited to nascent chromatin via its conserved TPR domain, which specifically interacts with H3.1 proteins unmethylated at K27 (H3.1K27me0) (*18*). In the absence of post-replicative chromatin maturation via H3.1K27me1, TSK is thought to either remain associated with chromatin post-replication or interact with it *de novo*. Genomic instability in *atxr5/6* mutants has been shown to be dependent on TSK activity (*18*), thus suggesting that the phenotypes observed in H3.1K27M-expressing plants may similarly rely on this protein.

To investigate a potential link between TSK and the H3.1K27M-associated phenotypes, we introduced the H3.1K27M transgene into a *tsk* mutant background (T-DNA insertional mutant SALK_034207). We found that elimination of TSK activity suppressed most of the growth abnormalities and early lethality associated with H3.1K27M expression in the Col-0 background (Fig. 4, A and B). Though the *tsk*/H3.1K27M plants remained smaller than Col-0, H3.1 WT, *tsk* and *tsk*/H3.1 WT plants, they were able to progress to the reproductive phase of growth, which was never observed when H3.1K27M was expressed in the Col-0 background. Flow cytometry analyses of *tsk*/H3.1K27M plants showed suppression of heterochromatin amplification as represented by the loss of broad peaks and confirmed by whole-genome sequencing (Fig. 4C and fig. S4, A and B). Chromocenter decondensation was also found to be suppressed in H3.1K27M-expressing plants lacking TSK (Fig. 4D). A possible mechanism of suppression is that H3K27me1 levels are restored to wild-type levels in a *tsk* mutant, however, this was ruled out by immunostaining and Western blot analyses (Fig. 4, D and E and fig. S4C). We also tested if suppression of the phenotypes associated with loss of H3.1K27me1 was due to lower H3.1K27M transgene expression in *tsk* mutants. We performed RT-qPCR and found no significant difference between Col-0 and *tsk* (fig. S4D). It is interesting to note that complete suppression of the genomic instability phenotypes in *tsk*/H3.1K27M plants does not lead to plants morphologically resembling Col-0 (Fig. 4, A and B). This is likely due to the loss of H3K27me3 in these plants caused by inhibition of PRC2 (Fig. 4F). Similar phenotypes were observed in the H3.1_prom_::H3.3K27M plants, where H3K27me3 is reduced, but not H3K27me1 (Fig. 2, E to H and fig. S4E).

**Figure 4.**
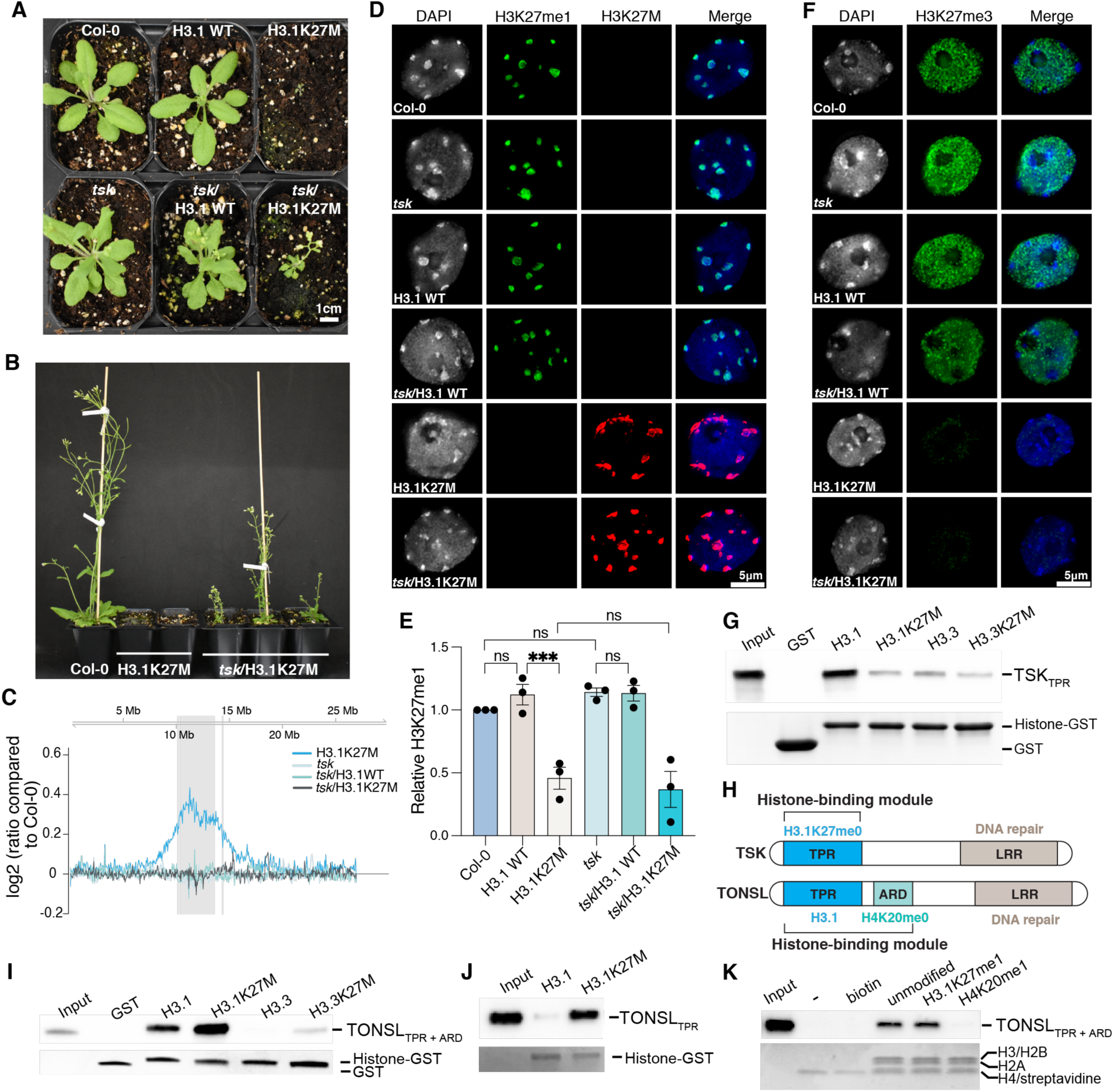
H3.1K27M causes genomic instability by inducing ectopic TSK activity. **A** and **B** Morphological phenotypes of Col-0, *tsk*, H3.1 WT, H3.1K27M, *tsk*/H3.1 WT and *tsk*/H3.1K27M plants. **C** Chromosomal view (Chr 5) of DNA-seq reads. The pericentromeric region is highlighted in gray. **D** Immunostaining of leaf nuclei with anti-H3K27M and anti-H3K27me1 antibodies. DNA is stained with DAPI. **E** Western blot quantification showing relative H3K27me1 levels in total histones extracted from leaves. One-way ANOVA with Tukey’s multiple comparison test: *** *p* < 0.001, ns = not significantly different. **F** Staining of leaf nuclei with an anti-H3K27me3 antibody and DAPI. **G** Peptide pull-down assay using the TPR domain of TSK (TSK_TPR_) and GST-tagged histone peptides (aa 1 to 58). **H** Domain architecture of plant and animal TSK/TONSL. TPR: Tetratricopeptide Repeats, ARD: Ankyrin Repeat Domain, LRR: Leucine-Rich Repeats. **I** and **J** Pull-down assay with TPR + ARD domains (TONSL_TPR + ARD_) (I) or the TONSL TPR domain only (TONSL_TPR_) (J) with GST-tagged histone peptides (aa 1 to 58). **K** Pull-down assay of TONSL_TPR + ARD_ with biotinylated recombinant nucleosomes.

Our findings suggest that H3.1K27M inhibits ATXR5/6 in a dominant-negative manner, which leads to an increase in K27me0 on wild-type H3.1 proteins. This results in chromatin binding of TSK outside of its normal spatial and temporal boundaries, and as a consequence, DNA damage. However, it is also possible that H3.1K27M increases the interaction of TSK with H3.1, thus leading to DNA damage at genomic regions where H3.1K27M is inserted. Analysis of the structure of the TSK_TPR_-H3.1 complex does not support this mechanism, as H3.1K27 is positioned within a polar pocket that makes it unfavorable for binding hydrophobic entities such as the sidechain of methionine (*18*). We verified this by performing *in vitro* binding assays and confirmed that the interaction of TSK with H3.1 is reduced when lysine 27 is replaced with methionine (Fig. 4G). We also conducted split-luciferase complementation assays and detected luminescence signals when TSK_TPR_-cLuc was coexpressed with H3.1 WT-nLuc, but not with H3.1K27M-nLuc (fig. S4, F and G). Altogether, these findings support a model where, in plants, genomic instability is not due to a direct interaction between H3.1K27M nucleosomes and TSK, but instead arises from increased levels of H3.1K27me0 leading to ectopic TSK activity.

To assess if the inhibitory effect of H3.1K27M and H3.1K27 methylation on TSK binding is conserved in the mammalian TONSL ortholog, we performed binding assays using the complete histone interaction module of TONSL (TPR and ARD domains) or the TPR domain only (Fig. 4H). Interestingly, we found that mouse TONSL_TPR+ARD_ and TONSL_TPR_ displayed increased binding to H3.1K27M compared to wild-type H3.1 (Fig. 4I-J), and that the interaction of TONSL_TPR+ARD_ with histones is restricted by H4K20me1, but not H3.1K27me1 (Fig. 4K). These results suggest that H3.1K27M insertion on chromatin leads to ectopic TSK/TONSL activity in both plants and animals, but via a different mechanism.

### H3.1K27M-expressing plants are hypersensitive to the loss of specific DNA repair proteins

H3.1K27M expression has similar effects on genomic stability as *atxr5/6* mutants. Our previous work showed that various phenotypes in *atxr5/6* mutants are either enhanced or suppressed by the loss of specific DNA repair pathways (*18*). For example, loss of RAD51 (involved in homologous recombination) or Pol θ (key player in theta-mediated end joining [TMEJ]) in *atxr5/6* mutants enhances the growth defects of these plants. This is likely due to an increased reliance on repair of damaged DNA resulting from higher levels of H3.1K27me0 and unregulated TSK activity. In Col-0, mutations in *RAD51* or *POLQ* (coding for POL θ) have no effects on plant growth (*18*). These observations suggest that plants expressing H3.1K27M may also be hypersensitive to the loss of DNA repair pathways due to high levels of DNA damage caused by the oncohistone. To test this, H3.1K27M was first introduced into heterozygous *rad51* mutants (T0 plants), as homozygous *rad51* mutants are sterile and therefore cannot be transformed by Agrobacterium using the flower dip method (*49*). Transformation of heterozygous *rad51* mutants with constructs expressing H3.1K27M or H3.1 WT resulted in 1% (1/95) and 25% homozygous *rad51* mutants (T1 plants), respectively (Fig. 5A). The near absence of homozygous *rad51* mutants in the T1 generation of H3.1K27M plants strongly suggests that homologous recombination is a major contributor to DNA repair and plant/cell viability in H3.1K27M-expressing plants. We could not replicate this strategy to assess the contribution of TMEJ via Pol θ in H3.1K27M plants, as *polq* mutants cannot be transformed via Agrobacterium-mediated flower dipping (*50*). As TMEJ relies on resected DNA as a substrate to initiate DNA repair (*51, 52*), we tested the effect of MRE11 in plants expressing H3.1K27M. MRE11 is a key component of the MRE11–RAD50–NBS1 (MRN) complex responsible for DNA end resection (*53*). Similarly to the experiment using *rad51*, we introduced H3.1K27M and H3.1 WT into heterozygous T0 plants since homozygous *mre11* mutants are sterile (*54*). We only recovered 2% (1/59) homozygous *mre11* mutants among the T1 progeny (Fig. 5B). In contrast, transformation of H3.1 WT resulted in 23% of the T1 transformants being homozygous *mre11* mutant, which is close to the expected 25% for a transgene with no effect on plant survival. This result suggests that DNA resection mediated by the MRN complex specifically contributes to the survival of H3.1K27M plants. To further explore if H3.1K27M-expressing plants respond like *atxr5/6* mutants to the absence of key DNA repair pathways, we tested non-homologous end joining (NHEJ) by directly transforming H3.1K27M or H3.1 WT into homozygous *ku80* mutants. The absence of Ku80 in plants expressing H3.1K27M did not enhance growth defects or suppress heterochromatin amplification, which parallels the phenotypes observed in *atxr5/6 ku80* triple mutants (Fig. 5C and fig. S5, A and B) (*18*). Finally, we assessed the impact of RAD17 in H3.1K27M-expressing plants. As observed in *atxr5/6 rad17* triple mutants (*18*), loss of RAD17 in H3.1K27M plants abolished heterochromatin amplification while having no measurable effects on plant growth (Fig. 5D and fig. S5, B and C). Overall, these results demonstrate that the phenotypes of H3.1K27M-expressing plants and *atxr5/6* mutants are modulated by the same DNA repair proteins, and that expression of the oncohistone in Arabidopsis generates a dependence on specific DNA repair pathways for survival.

**Figure 5.**
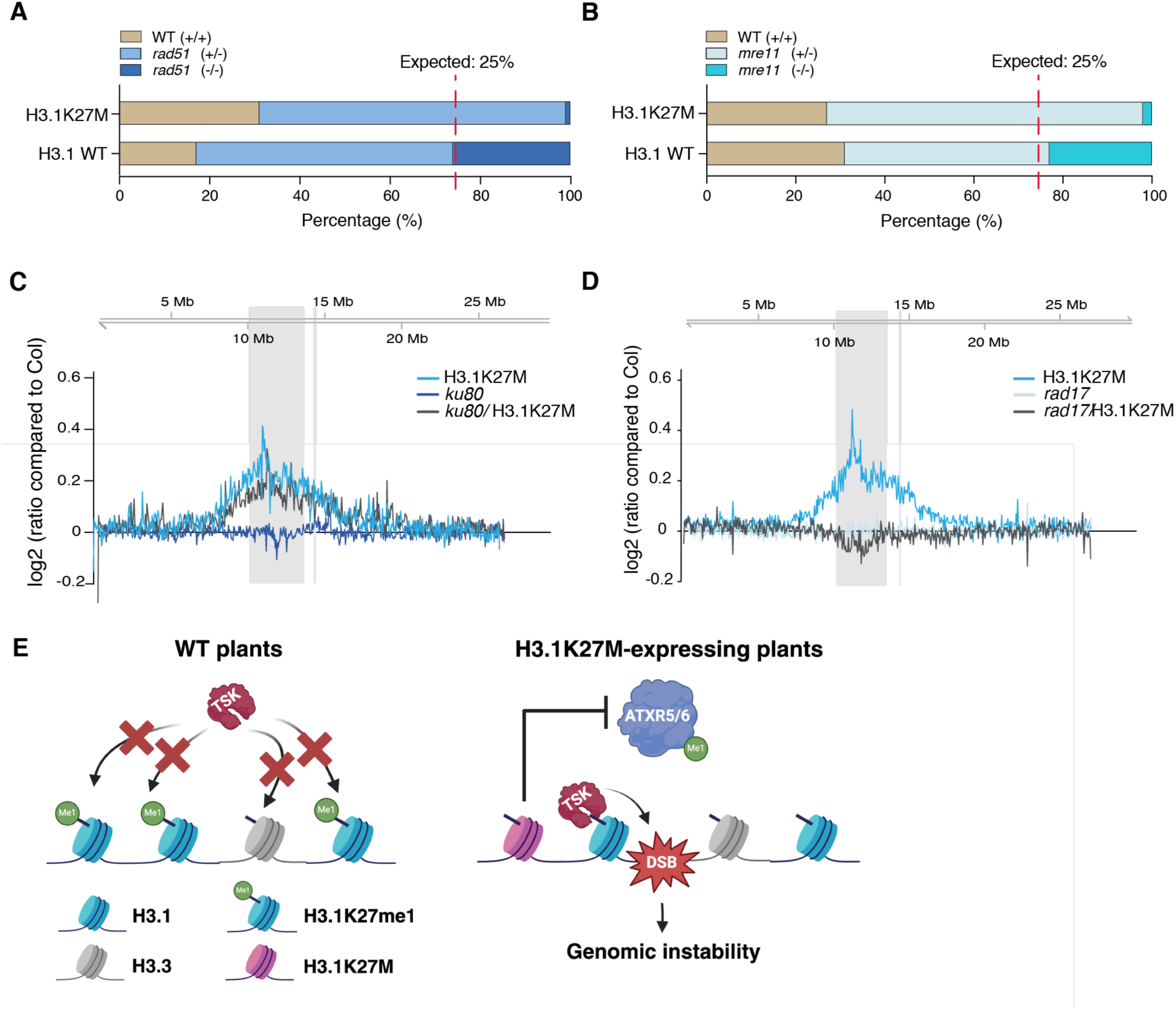
Differential effects of H3.1K27M expression in DNA repair mutants. **A** and **B** Genotypes of T1 transgenic plants resulting from transformation of H3.1 WT and H3.1K27M into *rad51* heterozygous (*rad51* [+/-]) (A) and *mre11* heterozygous (*mre11* [+/-]) (B) T0 plants. **C** and **D** Chromosomal view (Chr 5) of DNA-seq reads. The pericentromeric region is highlighted in gray. **E** Model depicting how H3.1K27M expression leads to genomic instability by increasing the levels of H3.1K27me0, which causes misregulation of TSK.

## Discussion

Genomic instability is associated with DMG-H3K27a and the H3K27M oncomutation (*29–33*). However, underlying mechanisms directly linking H3K27M to DNA damage in these cancers remains mostly unexplored. In this study, we showed using Arabidopsis as a model that expression of H3.1K27M replicates the genomic instability phenotypes observed in *atxr5/6* mutants, which are caused by misregulation of the TSK-H3.1 DNA repair pathway (*18*). Loss of H3.1K27me1 in *atxr5/6* mutants or H3.1K27M-expressing plants prevents post-replicative chromatin maturation, leaving H3.1 variants unmethylated. As H3.1K27me0 serves to recruit TSK to chromatin, a model emerges where ectopic binding of TSK to chromatin in H3.1K27M-expressing cells, or its retention on chromatin post-replication, induces DNA damage leading to genomic instability (Fig. 5E). More work will be needed to understand the mechanism by which TSK induces DNA damage when it is present on chromatin outside of its normal path of entry and exit during chromatin replication.

An important question is whether the findings of this study in plants translate to human cells, and if they can help us understand the etiology of DMG-H3K27a. In mammals, post-replicative chromatin maturation restricting TONSL activity has been shown to depend on SET8-catalyzed H4K20me1, which blocks the interaction between TONSL and chromatin (*23*). H4K20me0 is read by the TONSL ARD domain, which is not present in plant TSK orthologs (SET8 orthologs are also absent in plants) (*55*). Both TSK and TONSL contain the TPR domain that mediates the specific interaction with the H3.1 variant (*18*). TSK binding to H3.1 is inhibited by methylation of K27 in plants (*18*), but our *in vitro* results suggest that the TPR domain of TONSL is insensitive to mono-methylation at H3.1K27. This indicates that in contrast to plants, misregulation of TONSL activity in mammals may not occur as a result of genome-wide loss of H3K27 methylation due to PRC2 inhibition by H3.1K27M. However, and again in contrast to plant TSK, this study demonstrates that the TPR domain of TONSL displays enhanced binding to H3.1K27M compared to WT H3.1. Therefore, it is possible that H3.1K27M directly recruits TONSL to chromatin and form stable complexes that are not inhibited by the post-replicative chromatin maturation step mediated by SET8 via H4K20me1. This hypothesis will need to be tested in cell culture systems, but if confirmed, would support a model where H3.1K27M directly induces misregulation of TONSL activity, which may lead to DNA damage and genomic instability as in plants. Interestingly, our results suggest that the K27M mutation also increase the affinity of TONSL for the H3.3 variant compared to the non-mutated H3.3 protein. Therefore, there is a possibility that H3.3K27M expression may also lead to the misregulation of TONSL, with consequences for genome maintenance.

Finally, our findings are in line with other studies that point to diverse effects of H3K27M on the regulation of DNA repair pathways and genomic stability (*29–33*). These studies and our work suggest that H3.1K27M not only disrupts cellular identity via loss of H3K27me3, but also modulate the activity of DNA repair pathways to alter genomic integrity and induce tumorigenesis. Although this model underscores the incredible potency of H3K27M in disrupting key biological functions (i.e., epigenetic programming and genomic stability) required for cellular homeostasis, it also points to new therapeutic routes associated with the DNA damage response that could be applied to help manage or cure cancers characterized by this oncomutation.

## Methods

### Plant materials

Plants were grown at 22 °C under cool-white fluorescent lights (∼ 100 μmol m^−2^ s^−1^) in long-day conditions (16 h light/8 h dark). The T-DNA insertion mutants *atxr5/6* (At5g09790 /At5g24330, SALK_130607/SAIL_240_H01) (*25*), *tsk*/*bru1-4* (At3g18730, SALK_034207) (*56*), *rad51* (At5g20850, GK_134A01) (*49*), *ku80-7* (At1g48050, SALK_112921) (*57*), *rad17-2* (At5g66130, SALK_009384) (*58*) and *mre11* (At5g54260, SALK_028450) (*59*) are in the Col-0 ecotype background and were obtained from the Arabidopsis Biological Resource Center (Columbus, Ohio). Transgenic plants expressing H3.1 WT (HTR9), H3.1K27M (HTR9 K27M), and H3.1_prom_::H3.3K27M (HTR5 K27M driven by the HTR9 promoter) were made by transforming Col-0 plants or DNA repair mutants (*tsk*, *rad17*, *rad51*, *ku80* and *mre11*).

### Cloning

To generate transgenic expression constructs (H3.1 WT, H3.1K27M and H3.1_prom_::H3.3K27M), the coding sequences of HTR9 (*At5g10400*) and its promoter (1027 bp upstream of the start codon) were cloned into MoClo system level 0 plasmids pICH41308 and pICH41295, respectively. The H3.1K27M mutation was engineered by site-directed mutagenesis (QuikChange II XL, Agilent Technologies, Santa Clara, CA). The terminator of HSP18.2 (At5g59720, 248 bp downstream of stop codon) was cloned into pICH41276. The level 1 clones were seamlessly assembled using Type-II restriction enzyme BsaI in the order of Promoter-CDS-terminator into destination vector pICH47802. The level 1 clones were later assembled into level 2 final clones combined with level 1 Ole::RFP selection cassette using BpiI, which allows for Agrobacterium transformation and RFP selection of transformants in planta.

To generate TSK/TONSL-TPR and TONSL-TPR+ARD expression constructs, the coding sequences of the TPR domains of *A. thaliana* TSK (aa 1-525) and of mouse TONSL (aa 1-465), and the joint domains TPR+ARD of TONSL (aa 1-693) were cloned into the pET32a vector as previously described (*18*). As for the GST-fused peptides, the coding sequences of the N-terminal tails of *A. thaliana* H3.1 (aa 1-58) and H3.3 (aa 1-58) were fused with a C-terminal GST tag by cloning into pET28 as previously described (*18*). The H3.1K27M and H3.3K27M mutations were introduced by site-directed mutagenesis using the QuikChange II XL kit (Agilent Technologies). Briefly, mutagenic primers containing the K27M substitution were designed and used for PCR amplification of the plasmid template. The PCR product was then treated with DpnI to digest the parental DNA template, followed by transformation into into *E. coli* XL10-Gold ultracompetent cells (Agilent Technologies). The plasmids were sequenced to verify the introduction of the desired mutations.

For the histone methyltransferase assays, ATXR6 (residues 25-349) was cloned into pGEX-6P as previously described (*25*).

### Plant transformation

Plant transformations were done using the floral dip method (*60*). Briefly, 400 ml of Agrobacterium (strain GV310) liquid culture grown overnight at 28°C was spun down at 3,220 x g for 25 min and resuspended in 500 mL of transformation solution (5% sucrose and 0.02% Silwet L-77). Arabidopsis flowers were dipped into the bacterial solution for 30 seconds. T1 transgenic plants were selected with the seed specific RFP reporter (OLE1-RFP) with a SFA Light Base (Nightsea, Lexington, MA). Transformations into mutants were either done with homozygous plants (*rad17* and *ku80*) or heterozygous plants (*tsk*, *rad51* and *mre11*), due to low seed set or sterility (*49, 61, 62*). Transgenic *tsk*, *rad51* and *mre11* homozygous plants were identified in the T1 population.

### Survival assay

Seeds were germinated and grown on ½ MS plates at 22 °C under cool-white fluorescent lights (∼ 100 μmol m^−2^ s^−1^) in long-day conditions (16 h light/8 h dark) for 5 days, transferred to soil and continued growing under the same conditions. The number of surviving plants were counted on day 0, 4, 8, 12, 16, and 20 after transferring to the soil. The survival rate was calculated as the number of live plants divided by the number of total seeds for each genotype.

### Anthocyanin measurements

Anthocyanin content was determined using a modified version of a previously published protocol (*63*). 200 μL of extraction buffer (50% methanol and 5% acetic acid) was added to 50 mg of ground leaf tissue, followed by centrifugation. The supernatant was carefully decanted into a new tube. This step was repeated to ensure complete transfer of the supernatant. Absorbances were measured at 530 nm and 637 nm wavelengths. Anthocyanin concentration, expressed as Abs530 per gram fresh weight (g F.W.), was calculated using the formula [Abs530 - (0.25 × Abs657)] × 5.

### Comet Assay

DNA damage was assessed in three-week-old plants using a Comet Assay Kit (Trevigen). In brief, leaves were sectioned into small fragments with a razor blade in 500 μl of PBS buffer containing 20 mM EDTA and maintained on ice. The nuclei were then released into suspension, filtered through a 50 μm nylon mesh into a fresh tube, and subsequently mixed with an equal volume of low-melting-point agarose. The mixture was immediately spread onto CometSlides. Following a 1-hour lysis step at 4°C, the slides were immersed in 1× Tris-acetate electrophoresis buffer for 30 minutes to allow unwinding of the DNA before electrophoresis at 4°C for 10 minutes. Post-electrophoresis, the slides were stained with SYBR Gold for visualization of the nuclei. Imaging was conducted using a Nikon Eclipse Ni-E microscope with excitation at 488 nm. Quantitative analysis of DNA damage was performed using the OpenComet plugin for ImageJ (*64*).

### MMS genotoxic assay

Seeds were germinated and grown on vertically-oriented ½ MS plates with or without 100μg/ml methyl methanesulfonate (MMS) (Thermo Fisher Scientific, Bohemia, NY) under cool-white fluorescent lights (∼ 100 μmol m^−2^ s^−1^) in long-day conditions (16 h light/8 h dark). Root length measurements were done 10 days (No MMS) or 20 days (100μg/ml MMS) after germination, respectively. The relative root length was calculated by dividing the root length of individual seedlings grown in the presence of MMS by the average root length of plants of the same genotype grown on ½ MS plates.

### Somatic recombination assay

H3.1 WT and H3.1K27M transgenic plants were generated by transforming a previously described inverted repeat GUS reporter line (*37*). Experiments were performed at least three times in four-week-old F3 plants as previously described (*18*).

### RNA-seq

For each biological replicate, leaves from three plants cultivated in the same flat were combined for RNA extraction using the RNeasy Plant Mini Kit (Qiagen, Hilden, Germany). The integrity of RNA was confirmed with the Agilent 2100 Bioanalyzer Nano RNA Assay (Agilent, Santa Clara, CA). Sequencing libraries were prepared at the Yale Center for Genome Analysis (YCGA) with 200 ng of total RNA using oligo-dT beads for poly A selection. Sequencing (101bp paired-end) was performed on an Illumina NovaSeq X Plus flow cell. Fastp (version 0.21.0 with default parameters) was used to filter and trim paired-end reads (*65*) and reads with quality inferior to 20 were removed. STAR (version 2.7.2a) was used to align the data against the Arabidopsis TAIR10 (*66*) genome with a maximum of two mismatches (--outFilterMismatchNmax 2) (*67*). Biological replicates were analyzed for consistency by principal componenet analysis (PCA) using FactoMineR (*68*) The PCA was performed using all genes with an average TPM over ≥ 5 in all samples (fig. S6). For the analysis of the percentage of H3.1 gene transcripts (HTR1, HTR2, HTR3, HTR9, HTR13 and H3.1 transgene), the trimmed reads were mapped and quantified using Salmon (salmon v1.4.0, default parameters) (*69*). For the expression analysis of DNA repair response genes, we utilized a curated list of genes previously assembled by the Britt laboratory. This comprehensive list, accessible via the Britt lab’s website (http://brittlab.ucdavis.edu/plant-dna-repair-genes), includes key genes involved in plant DNA repair mechanisms (see Table S1 for the complete gene list). Paired-end fragments corresponding to TEs were determined with featureCounts (version 1.6.4) (*70*). Transposable elements (TEs) were defined as previously described (*71*). TEs were considered to be upregulated if they showed a ≥ 2-fold up-regulation as compared to Col in both biological replicates, and had a value of TPM ≥ 1. The heatmap was generated with the R heatmap.2 function from the ggplot2 package (*72*).

### Nuclei DAPI staining and Immunostaining

Two-week-old leaves were fixed in 3.7% formaldehyde in cold Tris buffer (10 mM Tris-HCl pH 7.5, 10 mM NaEDTA, 100 mM NaCl) for 20 minutes under vacuum. The formaldehyde solution was removed, and leaves were washed twice for 10 minutes in Tris buffer. The leaves were chopped using razor blades in 150 μl LB01 buffer (15 mM Tris-HCl pH7.5, 2 mM NaEDTA, 0.5 mM spermine-4HCl, 80 mM KCl, 20 mM NaCl and 0.1% Triton X-100), then filtered through a 30 μm mesh (Sysmex Partec, Gorlitz, Germany). 10 μl of lysate was added to 10 μl of sorting buffer (100 mM Tris-HCl pH 7.5, 50 mM KCl, 2mM MgCl_2_, 0.05% Tween-20 and 5% sucrose) and dried onto a coverslip for 30 minutes. Cold methanol was added onto each coverslip for 3 minutes. Methanol was aspirated and TBS-Tx (20 mM Tris pH 7.5, 100 mM NaCl, 0.1% Triton X-100) was added for 5 minutes. TBS-Tx was then removed. For DAPI staining only, the coverslips were immediately mounted onto slides with Vectashield mounting medium with DAPI (Vector Laboratories, Burlingame, CA). For immunostaining experiments, the following steps were performed: the coverslips were blocked by adding 75 μl of Abdil-Tx (2% BSA in TBS-Tx) to each coverslip for 30 minutes, followed by a 1 hour incubation with the appropriate primary antibody (H3K27me1; Active Motif #61015, H3K27me3, Millipore #07-449, H3K27M, Active Motif #61803) diluted in Abdil-Tx. The coverslips were then be washed with TBS-Tx. 30 μl of secondary antibody was added on each coverslip and incubated for 30 minutes followed by washing with TBS-Tx. Finally, the coverslips were mounted onto slides with Vectashield mounting medium with DAPI. Images (Z-series optical sections of 0.3 μm steps) were taken on a Nikon Eclipse Ni-E microscope with a 100X CFI PlanApo Lamda objective (Nikon, Minato City, Tokyo, Japan) equipped an Andor Clara camera. Images were deconvolved using the imageJ deconvolution plugin.

### ChIP-seq

ChIP was performed as described previously (*73*), with some modifications. Briefly, rosette leaves from 2-week-old plants were fixed for 15 min in 1% formaldehyde, followed by flash freezing. Approximately 0.1 g of ground tissue was added to 10 mL of extraction buffer 1 (0.4 M sucrose, 10 mM Tris–HCl, 10 mM MgCl_2_) and filtered through a 40 *µ*m mesh. Samples were centrifuge at 3,000*g* for 20 min. The pellets were resuspended in 1 mL of extraction buffer 2 (0.25 M sucrose, 10 mM Tris–HCl, 10 mM MgCl_2_, 1% Triton X-100) and centrifuged at 12,000*g* for 10 min. 400 *µ*L of extraction buffer 3 (1.7 M sucrose, 10 mM Tris-HCl (pH8.0), 0.15% Triton X-100) was used to resuspend the pellets. The homogenate was added to a fresh tube containing 400 *µ*L of extraction buffer 3, followed by centrifugation for 1 h at 16,000*g*. The pellets were resuspended in nuclei lysis buffer (50 mM Tris–HCl pH 8.0, 10 mM EDTA, 1% SDS), and chromatin was sheared using a Bioruptor 200 sonicator (20 times on a 30-s ON, 30-s OFF cycle). The supernatants were centrifuged at 16,000*g* for 5 min. ChIP dilution buffer (1.1% Triton X-100, 1.2 mM EDTA, 16.7 mM Tris–HCl pH 8.0, 167 mM NaCl) was added to the supernatant to obtain 10× volume. Antibodies (2 μL of H3K27me3 antibody (Millipore #07-449) or 2 μL of H3K27me1 antibody (ABclonal A22170) were added to 750 μL of diluted sample and incubated at 4°C overnight. An equal amount of drosophila chromatin (Active Motif #53083) was added to each sample to allow for quantitative comparisons of samples displaying very different amounts of H3K27 methylation marks (ChIP-Rx) (*74*). Immunoprecipitation was performed using protein A magnetic beads (New England BioLabs, Ipswich, MA). The beads were washed, resuspended in 200 *µ*L of elution buffer (1% SDS and 0.1-M NaHCO_3_) and incubated at 65°C for 15 min for elution. After uncrosslinking, samples were treated with proteinase K and purified using a ChIP DNA Clean and Concentrator kit (Zymo Research, Irvine, CA).

ChIP sequencing libraries were prepared at the YCGA with a TruSeq Library Prep Kit (Illumina, San Diego, CA). They were validated using an Agilent Bioanalyzer 2100 High sensitivity DNA assay, and quantified using the KAPA Library Quantification Kit for Illumina® Platforms kit. Sequencing was done on an Illumina NovaSeq 6000 using the S4 XP workflow. Fastp (version 0.21.0 with default parameters) was used to filter and trim paired-end reads (*65*). The reads with quality scores < 20 were removed. Duplicate reads were also removed using the Picard toolkit (https://broadinstitute.github.io/picard) (MarkDuplicates with REMOVE_DUPLICATES=true). To calculate the Rx scaling factor of each biological replicate, Drosophila-derived IP read counts were normalized to the number of input reads. Spike-in normalization was performed as previously described (*75, 76*). We used α = *r*/*Nd_IP* (*74*) to compute the scaling factor α for each replicate, with Nd_IP corresponding to the number of reads (in millions) aligning to the *D. melanogaster* genome in the IP and with *r* = 100 * *Nd_i / (Na_i + Nd_i*), where Nd_i and Na_i are the number of input reads (in millions) aligning to the *D. melanogaster* or *A. thaliana* genome, respectively. We generated bedgraph files with a bin size of 10 bp using deepTools (*77*) using Rx factors to scale each of the samples. The plot profile and heatmap in Fig. 2B were generated using deeptools. In order to generate the chromosomal representation in Figure 2C, featureCounts (version 1.6.4) was used to count the paired-end reads within 100-kb regions of the genome and scaled by adjusting the number of reads in each bin with Rx factors Consistency between biological replicates was confirmed by Pearson correlation using deepTools2 (fig. S7) (*77*).

### Histone protein extraction

Histone protein extraction was performed as previously described (*78*) with some modifications. For each sample, approximately 80 mg of plant material was ground with 1 ml of cold Nuclear Isolation Buffer (NIB) (*78*) and transferred to a 2-ml Dounce homogenizer. The plant homogenate was then processed with a loose pestle in the Dounce homogenizer for 50 strokes on ice. The lysate was filtered through two layers of Miracloth into new tubes and centrifuged. The supernatant was discarded, and the pellet was washed three times with NIB. The nuclear pellet was then resuspended in 1 ml of 0.4 N H2SO4. This mixture was transferred to a 2-ml Dounce homogenizer and homogenized with 40 strokes on ice to disrupt the nuclei. After centrifugation, the supernatant was transferred to a fresh tube and mixed with 264 μl of trichloroacetic acid dropwise to precipitate the proteins. The mixture was incubated on ice for 3 hours to ensure complete precipitation. Following incubation, the histone pellet was centrifuged at 14,000 rpm for 15 minutes at 4°C. The supernatant was carefully removed, and the pellet was washed with 500 μl of ice-cold acetone to remove any remaining acid and impurities. The acetone wash step was repeated once more to ensure thorough cleaning. The pellet was then air-dried at room temperature until no visible moisture remained. Finally, the dried histone pellet was resuspended in 25 μl of histone storage buffer (10 mM Tris-HCl, pH 7.5, 0.1 mM EDTA) for storage at -80°C.

### Western blotting

Total histone samples were subjected to Western blot analysis. Laemmli sample buffer (Bio-Rad, Hercules, CA) was added to each total histone sample, incubated at 95°C for 3 minutes, and loaded onto a 4%-20% gradient SDS-PAGE gel. Following electrophoresis, proteins were transferred onto membranes using a semi-dry blotter in Tris-glycine transfer buffer containing 20% methanol. The membranes were then blocked with 5% non-fat dry milk in TBST (20 mM Tris-HCl, pH 7.6, 150 mM NaCl, 0.1% Tween-20) for 1 hour at room temperature to prevent non-specific binding. The membranes were then probed with H3K27me1(Active Motif: 61015) or H3K27me3 (Millipore: 07-449) for 1 hour, followed by incubation with a secondary HRP-labeled antibody (Sigma) at a 1:10 000 dilution for 1 hour at room temperature. Blots were visualized using the Bio-Rad Clarity Western ECL Substrate and imaged with a Bio-Rad ChemiDoc MP Imaging System. Relative quantification of Western blot bands was performed with Image J.

### Nucleosome assembly

Recombinant nucleosome arrays for histone methyltransferase assays were assembled as described previously (*79*). In short, histones were expressed in *E. coli* and purified from inclusion bodies. Histone octamers consisting of Arabidopsis H2A.13 (At3g20670), H2B (At3g45980), and Xenopus H4 along with either Xenopus H3 fused to a C-terminal Strep tag (H3-CT Strep) or Arabidopsis H3.1 (At1g09200) with K27A or K27M mutations were reconstituted by dialysis into refolding buffer (10 mM Tris pH 8, 1 mM EDTA, 5 mM b-mercaptoethanol, 2 M NaCl ) and purified by size exclusion chromatography on a Superdex 200 gel filtration column (GE Healthcare). To form nucleosome arrays, histone octamers were assembled onto a plasmid containing 12 repeats of the 601 nucleosome positioning sequence connected via a 30-bp linker using gradient dialysis from 2 M to 0.4 M NaCl followed by a step dialysis into TE.

### Histone methyltransferase assay

Histone methyltransferase assays were performed as previously described (*80*). Briefly, 0.5 ug of GST-ATXR6 and 0.5 ug of H3-CT-Strep nucleosomes were added to a total reaction volume of 25 ul. We either added no inhibitor, or 0.5 ug of K27A or K27M nucleosomes. The reactions were incubated for 1 h at 30 °C. SDS-PAGE sample buffer was added to each tube followed by heating to 95°C for 5 min. Samples were resolved by SDS-PAGE (15% gels) and transfered to a PVDF membrane. Coomassie stain solution (45% methanol, 10% acetic acid, 0.25% Coomassie Brilliant Blue R) was used to stain the membrane followed by destaining (45% methanol, 10% acetic acid). Membranes were air-dried, sprayed with EN^3^HANCE (Perkin Elmer) and exposed to autoradiography film overnight at -80 °C. Films were developed and bands were quantified using the software ImageJ.

### Flow cytometry

Rosette leaves from two-week-old plants were finely chopped in 0.2 – 0.5 ml Galbraith buffer (45 mM MgCl2, 20 mM MOPS, 30 mM sodium citrate, 0.1% Triton X-100, 40 μg/ml RNase A) and filtered through a 30 μm mesh (Sysmex Partec). Isolated nuclei were stained by adding 20 μg/ml propidium iodide (Sigma) to each sample, followed by vortexing. The samples were analyzed using BD FACSAria II sorter (Becton Dickinson, Franklin Lakes, NJ). FlowJo 10.9.0 (Tree Star, Ashland, Oregon) was used to generate profiles and for quantification (nuclei counts and rCV values). Each biological replicate represents one plant.

### DNA-seq

Genomic DNA was extracted as previously described (*81*). For each biological replicate, 5 plants were pooled to obtain approximately 40 mg of leaf tissue. DNA sequencing libraries were prepared at the YCGA. Genomic DNA was sonicated to an average fragment size of 350 bp using a Covaris E220 instrument (Covaris, Woburn, MA). The libraries were generated using the xGen Prism library prep kit for NGS (Integrated DNA Technologies, Coralville, IA). Paired-end 150 bp sequencing was performed on an Illumina NovaSeq 6000 using the S4 XP workflow (Illumina, San Diego, CA). Raw FASTQ data were filtered and trimmed using the fastp tool (--length_required 20, -- qualified_quality_phred 20) (*65*). The filtered sequences were then mapped to the genome (TAIR10) using BWA-MEM default parameters (https://doi.org/10.48550/arXiv.1303.3997). Mapped reads were sorted, duplicates were removed and indexed using SAMtools (*82*). Biological replicates were analyzed for consistency with deepTools2 (fig. S8) (*77*). The program featurecount (version 2.0.6) (*70*) was used to count the paired-end fragments present in each 50-kb region of the *A. thaliana* genome as previously described (*18*). The log2 ratio was centered on the average ratio of any two compared libraries normalized to the first 5 Mbp of chromosome 1. Plot profiles were generated using R (version 4.3.2) (*83*) and Gviz (*84*).

### Protein expression

Plant histones-GST fusion proteins and AtTSK-TPR were expressed in Rosetta (DE3) E. coli (Sigma, St. Louis, MO). The bacterial cultures were grown in Luria-Bertani (LB) medium, and expression of the fusion proteins were induced with 1 mM isopropyl β-D-1-thiogalactopyranoside (IPTG) at room temperature. Human histones-GST fusion proteins, mouse TONSL-TPR, mouse TONSL-TPR+ARD were expressed in ArcticExpress (DE3) E. coli (Agilent, Santa Clara, CA). The expression of human histones-GST fusion proteins, TONSL-TPR and TONSL-TPR+ARD were expressed at 16 ℃ for 24 hours and induced with IPTG (1mM) at OD_600nm_= 0.4.

To purify the histone-GST fusion proteins, harvested cell pellets were resuspended in PBS (1X Phosphate-Buffered Saline: 137 mM NaCl, 10 mM phosphate, 2.7 mM KCl, pH 7.4) supplemented with 1 mM phenylmethylsulfonyl fluoride (PMSF) prior to lysis by sonication and clarification by centrifugation. The clear lysate was then applied to a Glutathione Sepharose 4B affinity column. After thorough washing with PBS to remove unbound material, the fusion proteins were eluted with Elution Buffer (EB: 50 mM Tris-HCl, 50 mM NaCl, 30 mM reduced L-Glutathione, 10% glycerol, pH 8.0). The eluted proteins were then concentrated using Amicon® Ultra Centrifugal Filter Units (MilliporeSigma) with a 10 kDa molecular weight cutoff. The elution buffer was replaced with storage buffer (50 mM Tris-HCl, 50 mM NaCl, 10% glycerol, pH 8.0) by centrifuging at 4,000 x g for 20 minutes. The concentrated and purified proteins were aliquoted and preserved at -80°C for long-term storage.

ATXR6 expression in *E.coli* and purification was described in detail in a previous publication (*25*). The purification of AtTSK-TPR, TONSL-TPR and TONSL-TPR+ARD (containing an N-terminal Trx-His-S tag and a C-terminal His tag) was performed as previously described with minor modifications (*18*). Briefly, the cell pellets were resuspended in NPI-10 buffer (50 mM NaH_2_PO_4_, 300 mM NaCl, 10 mM imidazole, pH 8.0) containing 1 mM PMSF, and sonicated for 2.5 min (30 s pulse, 1 min break, 5 cycles). After centrifugation, the supernatant was transferred into a new 50 ml Falcon tube and added with washed agarose Ni-NTA His beads. The expressed proteins were trapped after passing through the Ni-NTA agarose column. The column was then washed by 7.5 ml NPI-20 (50 mM NaH2PO4, 300 mM NaCl, 20 mM imidazole, pH 8.0) twice, and eluted with 10 ml NPI-250 buffer (50 mM NaH2PO4, 300 mM NaCl, 250 mM imidazole, pH 8.0). The eluted proteins were further concentrated using Amicon Ultra Centrifugal Filter (50 kDa) to 1.5 ml for size exclusion chromatography (SEC). The concentrated proteins were injected onto SEC and eluted using a SEC buffer (25mM Hepes, 200mM NaCl, pH 7.5). Peak fractions from the S200 column were pooled, concentrated to 1-2 mg ml^−1^, flash frozen under liquid nitrogen and stored at −80 °C.

### Histone binding assays

For the TSK_TPR_ -histone binding assays, 2 μg of AtTSK were combined with either 2 μg of GST or GST-tagged histone tails (aa 1 to 58 of plant H3.1) in 400 μl of binding buffer (25 mM Tris-HCl, 250 mM NaCl, 0.05% NP-40, pH 8.0). The mixture was incubated overnight at 4°C with rotation. Subsequently, 15 μl of pre-washed Glutathione Sepharose 4B agarose beads (Cytiva, Marlborough, MA) were added to each sample and incubated for 30 minutes to pull-down the GST-tagged proteins. The beads were then washed four times for 5 minutes with 1 mL of binding buffer, while rotating at 4°C. The final wash was performed with binding buffer containing 150 mM NaCl. The proteins were eluted with 15 μl of 2× SDS loading buffer and denatured by boiling at 95°C for 5 minutes. Samples were then resolved on a 10% SDS-PAGE gel. The lower section of the gel underwent Coomassie staining to assess the GST and GST-tagged proteins, while the upper section was analyzed by Western blot using an anti-His antibody (Sigma; H1029). The TONSL_TPR +ARD_ and TONSL_TPR_ binding assays were performed as described above with the following modifications. The binding buffer consisted of 50 mM Tris-HCl pH8.0, 300 mM NaCl, 5% glycerol, 0.25% NP-40, 0.2 mM EDTA, 1 mM DTT. The binding reaction was incubated for 2 hours at 4°C. The samples were resolved on a 4-20% gradient gel. Each pull-down experiment was repeated a minimum of three times.

### Nucleosome binding assay

The nucleosome binding assays were performed as described for the histone binding assays with TONSL_TPR +ARD_. The Recombinant human biotinylated nucleosomes were obtained from EpiCypher (Durham, NC).

### Split-luciferase complementation assays

The TPR domain (aa 1-524) of TSK, fused with a nuclear localization signal (NLS), and the NLS alone, were cloned into the Gateway destination vector pGWB-cLUC (Addgene Plasmid #174051). Histones H3.1 and H3.1K27M (aa 1-136) were inserted into the Gateway destination vector pGWB-nLUC (Addgene Plasmid #174050). Agrobacterium tumefaciens GV3101 strains harboring these constructs were introduced into the leaves of 4-week-old Nicotiana benthamiana plants. For co-infiltration experiments, Agrobacterium cultures adjusted to cell densities of 0.1–0.5 at OD600 were mixed in equal volumes. To inhibit gene silencing, the tomato bushy stunt virus (TBSV) p19 silencing suppressor, encoded within the pDGB3alpha2_35S:P19:Tnos vector (Addgene Plasmid #68214), was also included in the infiltration mixture. The Agrobacterium inoculum was prepared by resuspension in infiltration buffer (10 mM MgCl2, 10 mM MES, pH 5.6, and 150 μM acetosyringone) and incubated at room temperature for 2 hours. The inoculum was then gently infiltrated into the abaxial side of the leaves using a 1 ml needleless syringe. At 48-72 hours post-infiltration, a 1 mM luciferin solution was applied to the leaves. Before imaging, detached leaves were kept in darkness for 10 minutes to reduce chlorophyll fluorescence interference. Luminescence was captured with a charge-coupled device (CCD) camera. Image analysis and quantification were performed using ImageJ software.

### RT-qPCR

RNA was isolated from two-week-old leaf tissue using TRIzol (Invitrogen, Carlsbad, CA). RNA samples were treated with RQ1 RNase-free DNase (Promega, Madison, WI) at 37°C for 30 min. 750 ng of total RNA was used to produce cDNA with iScript cDNA Synthesis Kit (Bio-Rad, Hercules, CA). Real-time PCR was done using KAPA SYBR FAST qPCR Master Mix (2X) Kit (Kapa Biosystems, Wilmington, MA) on a CFX96 Real-Time PCR Detection System (Bio-Rad, Hercules, CA). Relative quantities were calculated using the Ct method with *ACTIN7* (*At5g09810*) as the normalizer (*85*). At least three biological replicates were performed for each experiment.

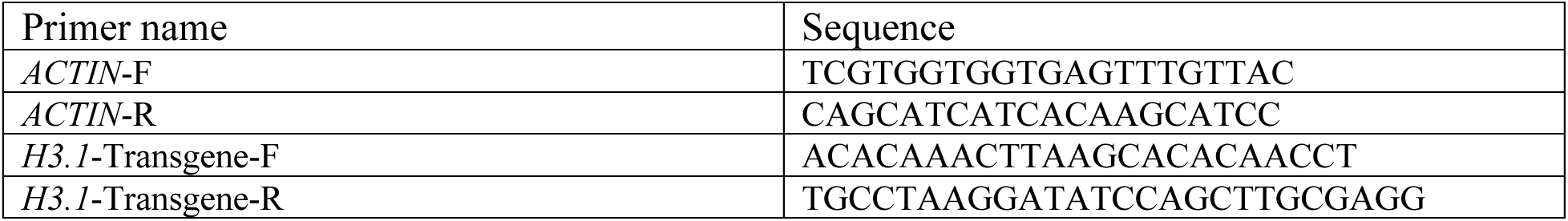

### Plant genotyping

Leaves from 2 or 3-week-old plants were homogenized in 500 μl DNA extraction buffer (200 mM Tris-HCl (pH 8.0), 250 mM NaCl, 25 mM EDTA and 1% SDS) and 50 μl phenol:chloroform:isoamyl alcohol (25:24:1). Each sample was centrifuged for 10 min at 15,000 rpm. 350 μl of the aqueous layer was transferred to a 1.5 ml tube containing 350 μl isopropanol. Samples were then mixed, incubated at room temperature for 15 min and then spun down at 15,000 rpm for 10 min. The supernatant was removed to secure the pellets. 400 μl 70% ethanol was used to wash the pellets. After centrifugation at 15,000 rpm for 5 min, the ethanol was removed and the pellets were dissolved in 50 μl water. Genotypes of individual T-DNA insertion T1 transgenic plants were determined by running two sets of genotyping PCR using GoTaq DNA polymerase (Promega, Madison, WI). WT primer sets amplify the original sequence of the given genes, while the mutant primer sets amplify the mutated sequence with T-DNA insert.

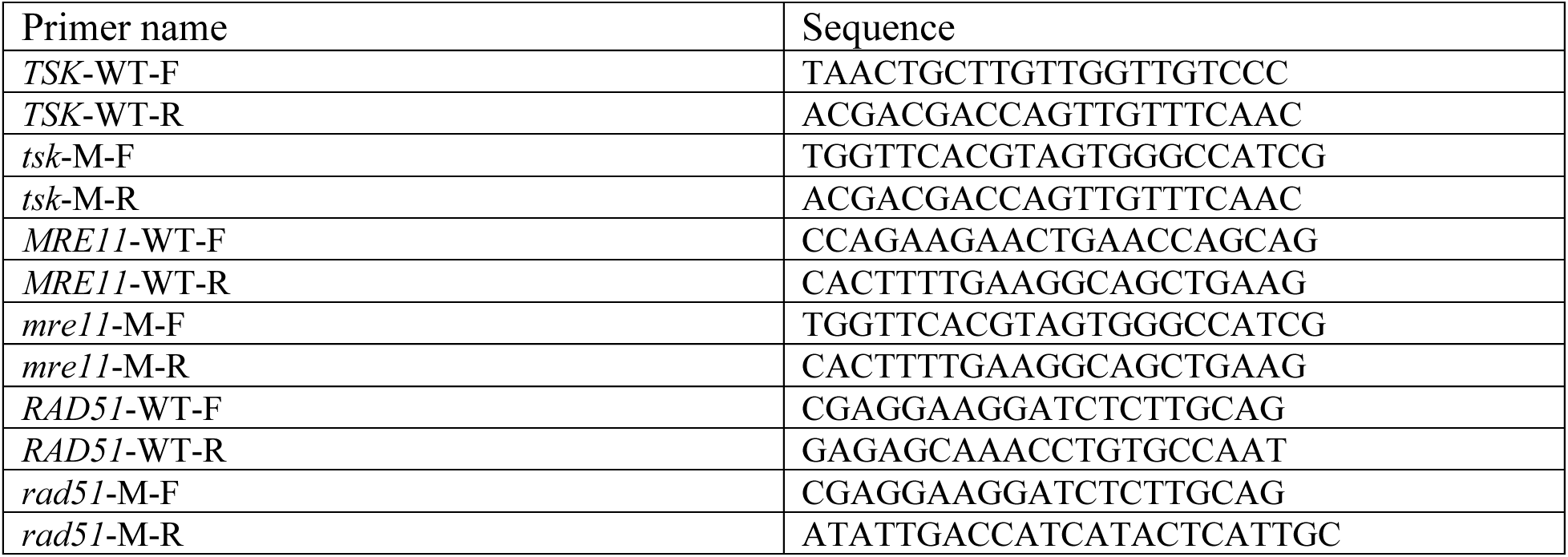

## Data availability

Sequencing data (RNA-seq, ChIP-seq and DNA-seq datasets) generated for this study are available from the SRA under accession numbers PRJN1121202. Additional materials reported in this study are available upon request from the corresponding author.

## Acknowledgments

We thank Ling Xu from Yale University for help with protein purification. We also thank members of the Jacob Laboratory for comments and discussion. This project was made possible by a grant (R35GM128661) from the National Institutes of Health (NIH) and a Yale Cancer Center Pilot Award to Y.J. Work in the Voigt lab was supported by the Wellcome Trust ([104175/Z/14/Z], Sir Henry Dale Fellowship (to P.V.) and the UK Biotechnology and Biological Sciences Research Council (BBS/E/B/000C0421). Support for this work in the lab of J.C.v.W. came from NIH grant R01AG078926. Research reported in this publication was also supported by the National Institute of General Medical Sciences of the NIH under Award Number 1S10OD030363-01A1 to the Yale Center for Genome Analysis.

## Author contributions

Y.J. supervised the work, and conceptualized the study and the experiments with Y.C.H. and W.Y. All the experiments were performed by W.Y., Y.C.H., and C.L. The genomic analyses were done by A.P. and W.Y. The nucleosomes arrays were generated by D.V. and P.V. Funding for the project was secured by Y.J., J.v.W., and P.V. The manuscript was written by Y.J. and C.L, with contributions from W.Y. and Y.C.H.

## Ethics declarations

The authors declare no competing interests.

