## Supplemental data for "H3.1K27M-induced misregulation of the TSK/TONSL-H3.1 pathway causes genomic instability"

**This PDF file includes:**

Figs. S1 to S8
Legends for tables S1 to S2

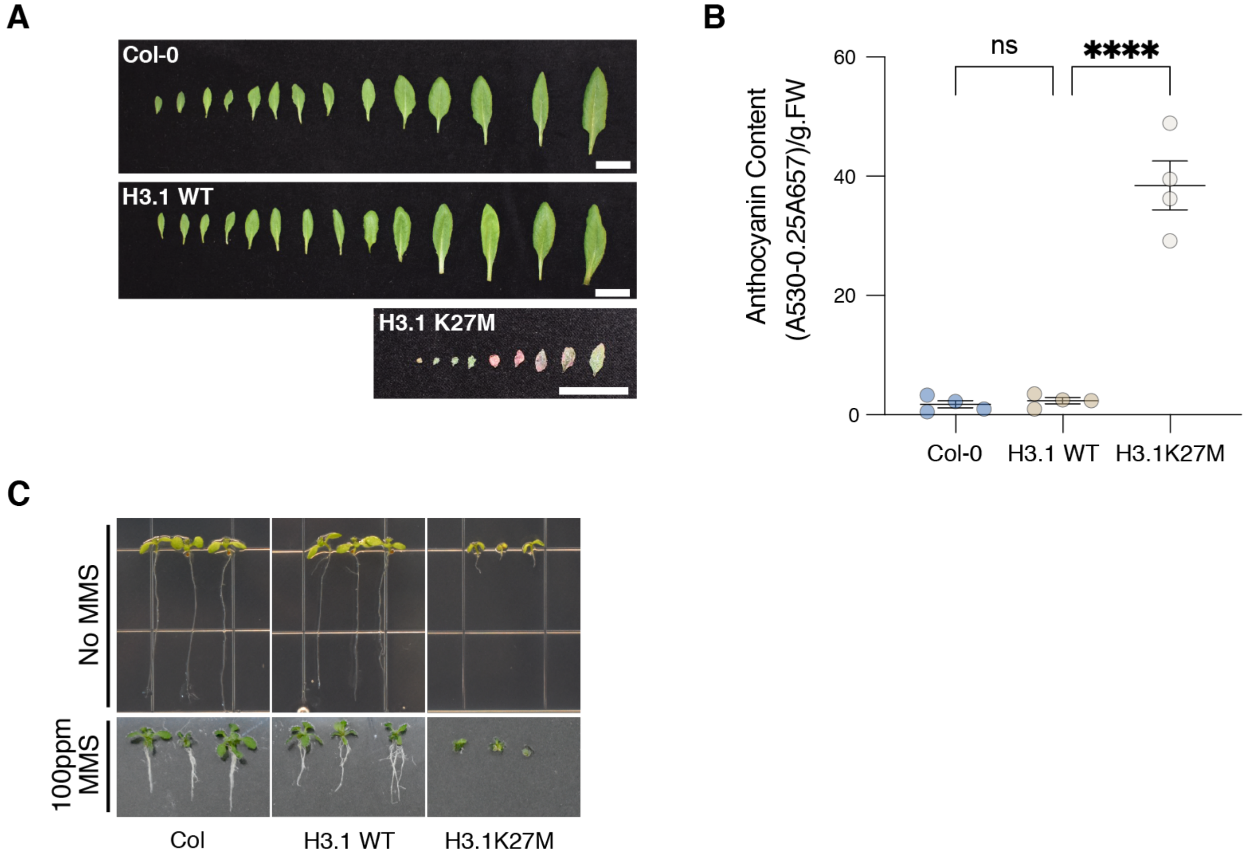


**Fig. S1. H3.1K27M expression increases anthocyanin content and sensitivity to MMS. A** Rosette leaves of Col-0, H3.1 WT and H3.1K27M plants. Scale bar = 1 cm. **B** Quantification of total anthocyanin contents in leaves from Col-0, H3.1 WT and H3.1K27M plants. Each dot represents and individual T1 plant. Horizontal bars indicate the mean. SEM is shown. One-way ANOVA with Tukey's multiple comparison test: **** $p < 0.0001$, ns = not significantly different. **C** Seedlings grown on vertically-oriented plates with and without MMS.

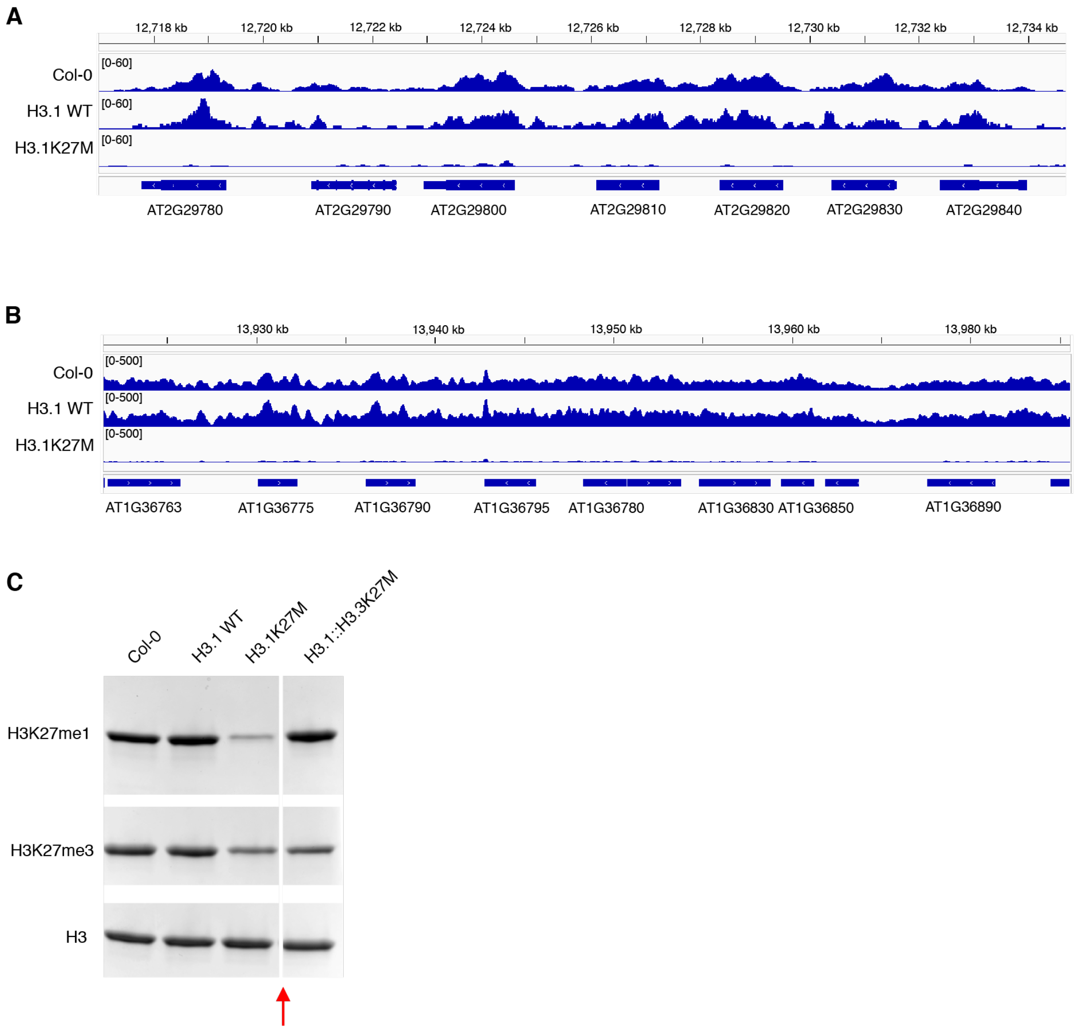


**Fig. S2. H3K27 methylation is decreased in H3.1K27M plants. A** and **B** Genome browser snapshots of ChIP-seq data showing normalized H3K27me3 over a euchromatic region of chr 2 (A) and H3K27me1 over a heterochromatic region of chr 1 (B). **C** Representative Western blot showing relative H3K27me1and H3K27me3 levels in total histones extracted from leaves. The red arrow indicates a gel lane that was removed.

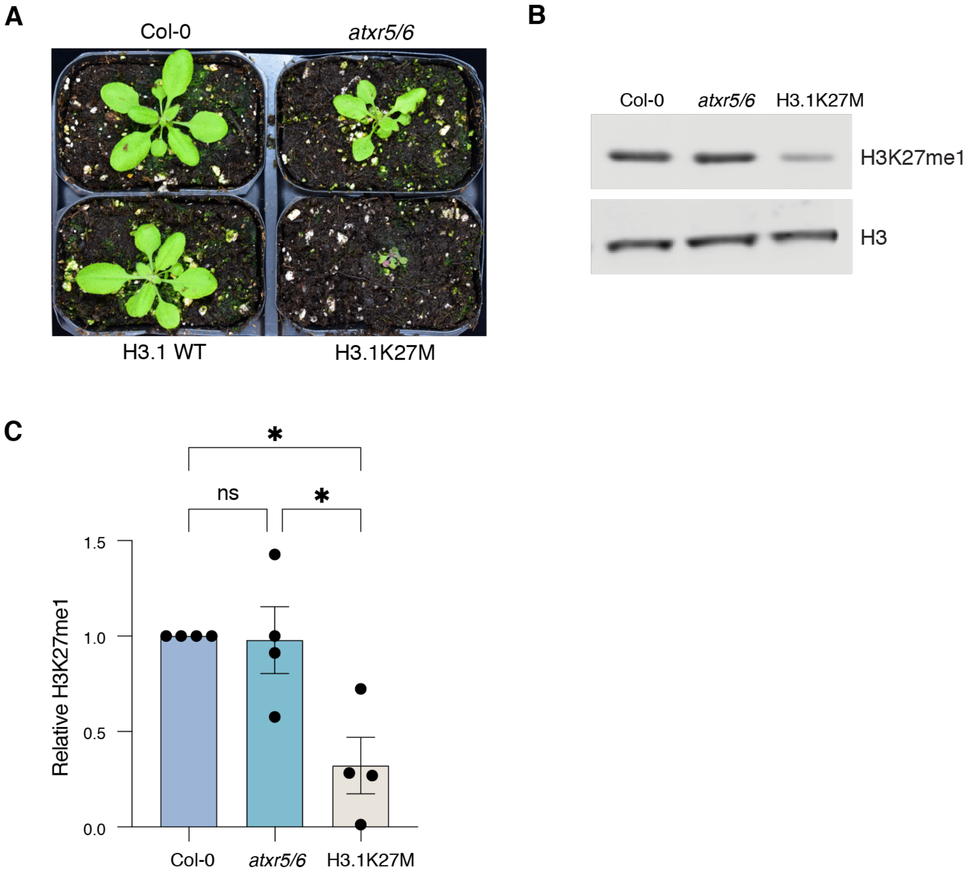


**Fig. S3. H3K27me1 levels are lower in H3.1K27M plants compared to *atxr5/6* mutants. A** Morphological phenotypes of Col-0, *atxr5/6*, H3.1 WT and H3.1K27M plants. **B** Representative Western blot showing H3K27me1 in total histones extracted from 2-week-old leaves. **C** Western blot quantification of H3K27me1 in Col, *atxr5/6* and H3.1K27M from four independent experiments. Horizontal bars indicate the mean. SEM is shown. One-way ANOVA with Tukey's multiple comparison test: * $p < 0.05$, ns = not significantly different.

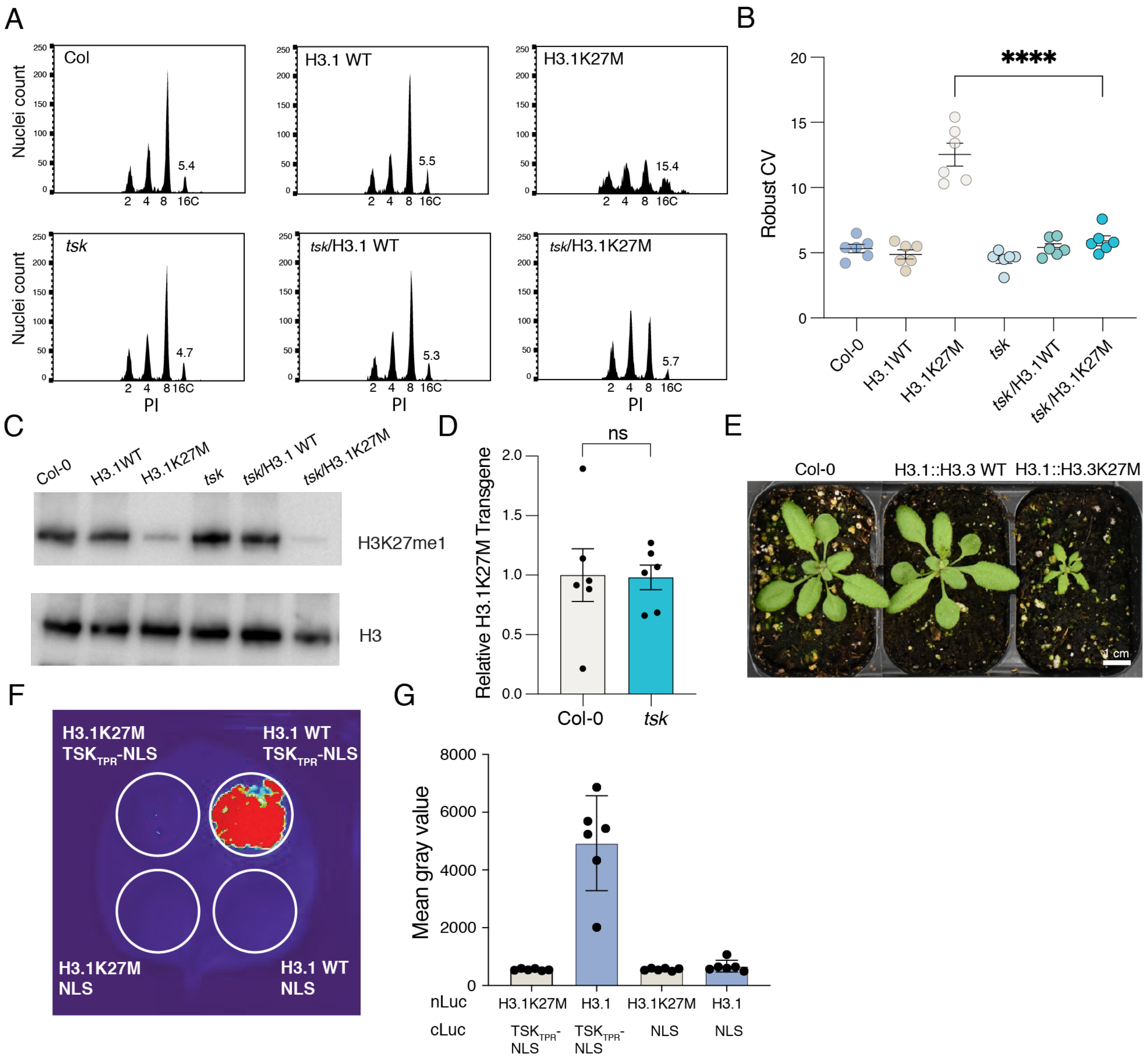


**Fig. S4. Genomic instability caused by H3.1K27M is TSK-dependent. A** Flow cytometry profiles of leaf nuclei stained with PI. Ploidy of the nuclei are shown below the peaks. The numbers above the 16C peaks indicate the robust CV. **B** Robust CV quantification of 16C peaks. Each dot represents an individual plant. The mean and SEM are shown. One-way ANOVA with Tukey's multiple comparison test: **** $p < 0.0001$. **C** Representative Western blot showing levels of H3K27me1 in total histones extracted from leaves. **D** RT-qPCR analysis of the H3.1K27M transgene in Col-0 and *tsk* plants. Each dot represents one T1 plant. Unpaired *t*-test: ns = not significantly different. **E** Morphological phenotypes of Col-0, H3.1::H3.3 WT and H3.1::H3.3K27M plants. **F** Split-LUC imaging assay. Agrobacterium carrying H3.1 WT or H3.1K27M and $TSK_{TPR}$-NLS fused to the N- and C-terminal domain of luciferase, respectively, was infiltrated in different locations on an *N. benthamiana* leaf. The NLS fused to the C-terminal domain of luciferase was used as a negative control. **G** Quantification of assay shown in (F) using a microplate luminescence reader.

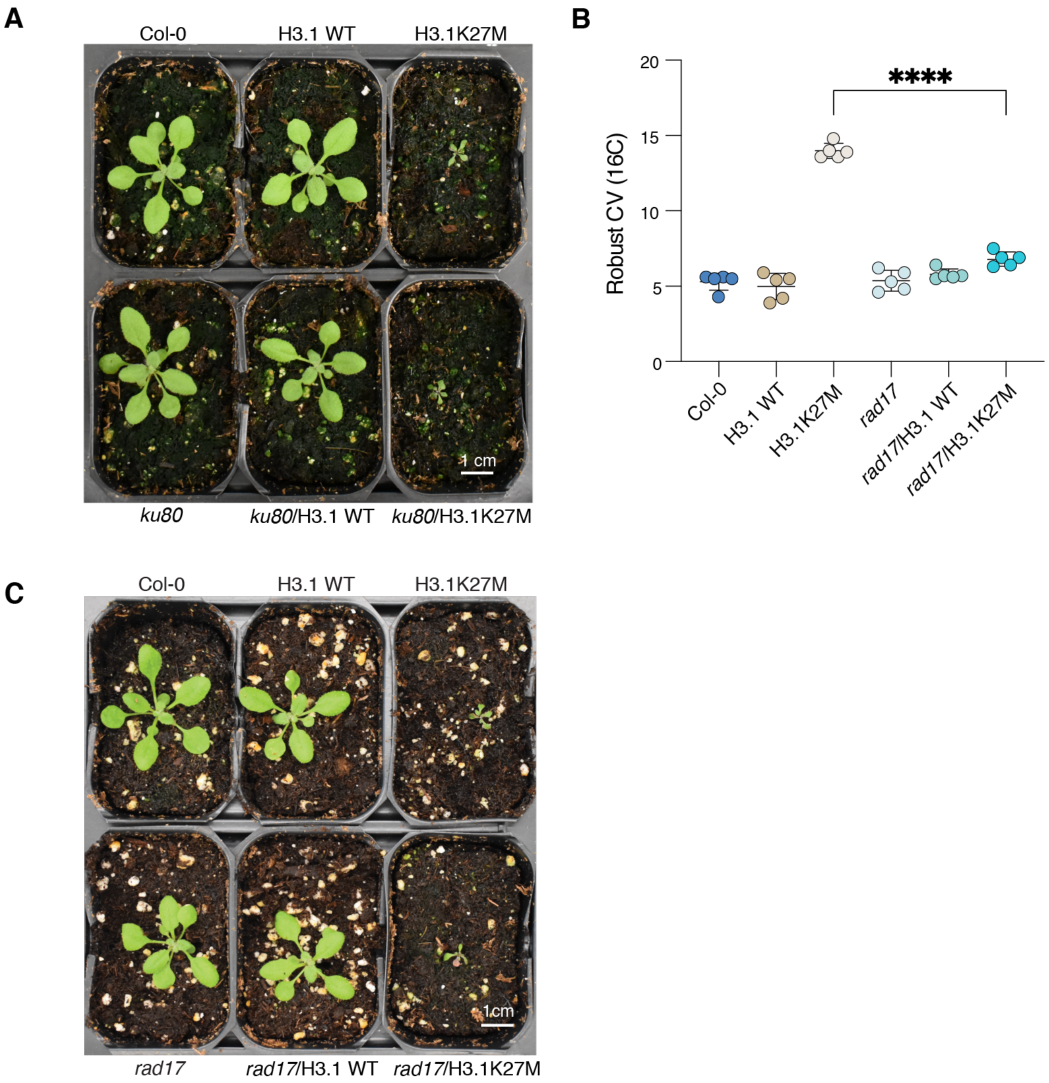


**Fig. S5. Effects of inactivation of *ku80* and *rad17* on H3.1K27M plants**. **A** Morphological phenotypes of *ku80* H3.1K27M-expressing plants. **B** Robust CV quantification of 16C peaks of flow cytometry profiles. Each dot represents an individual plant. The mean and SEM are shown. One-way ANOVA with Tukey's multiple comparison test: **** $p < 0.0001$. **C** Morphological phenotypes of *rad17* H3.1K27M-expressing plants.

**A**

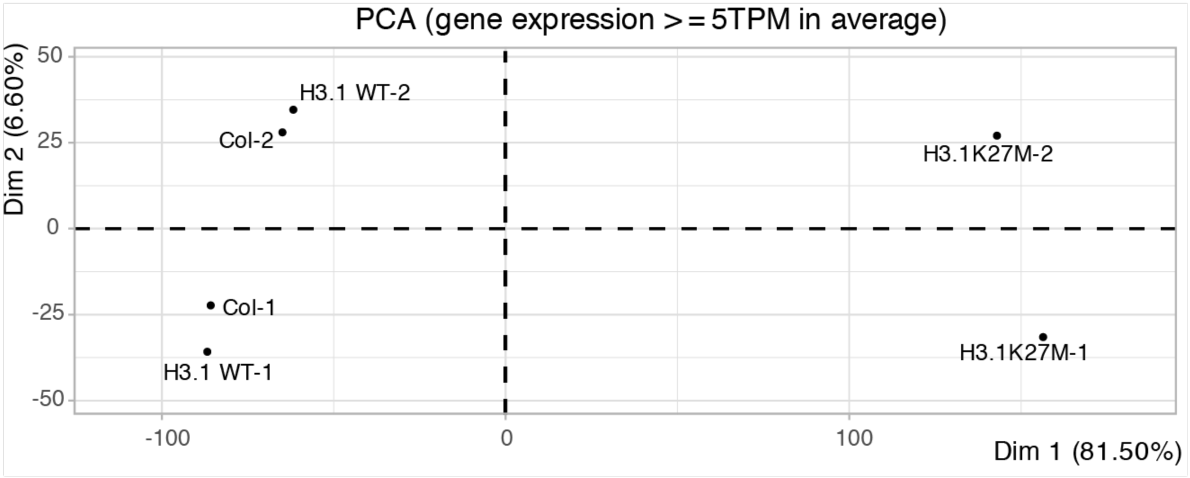


**B**

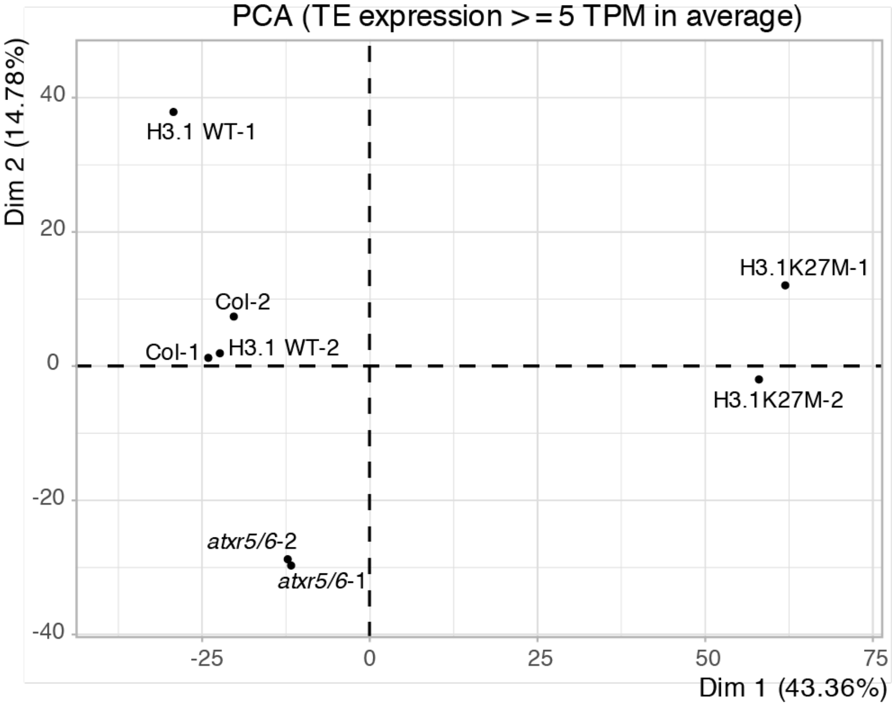


**Fig. S6. PCA scatter plot of RNA-seq samples. A** PCA scatter plot for genes > 5 TPM**. B** Plots for TEs > 5 TPM. Each dot represents a biological replicate of RNA-seq samples.

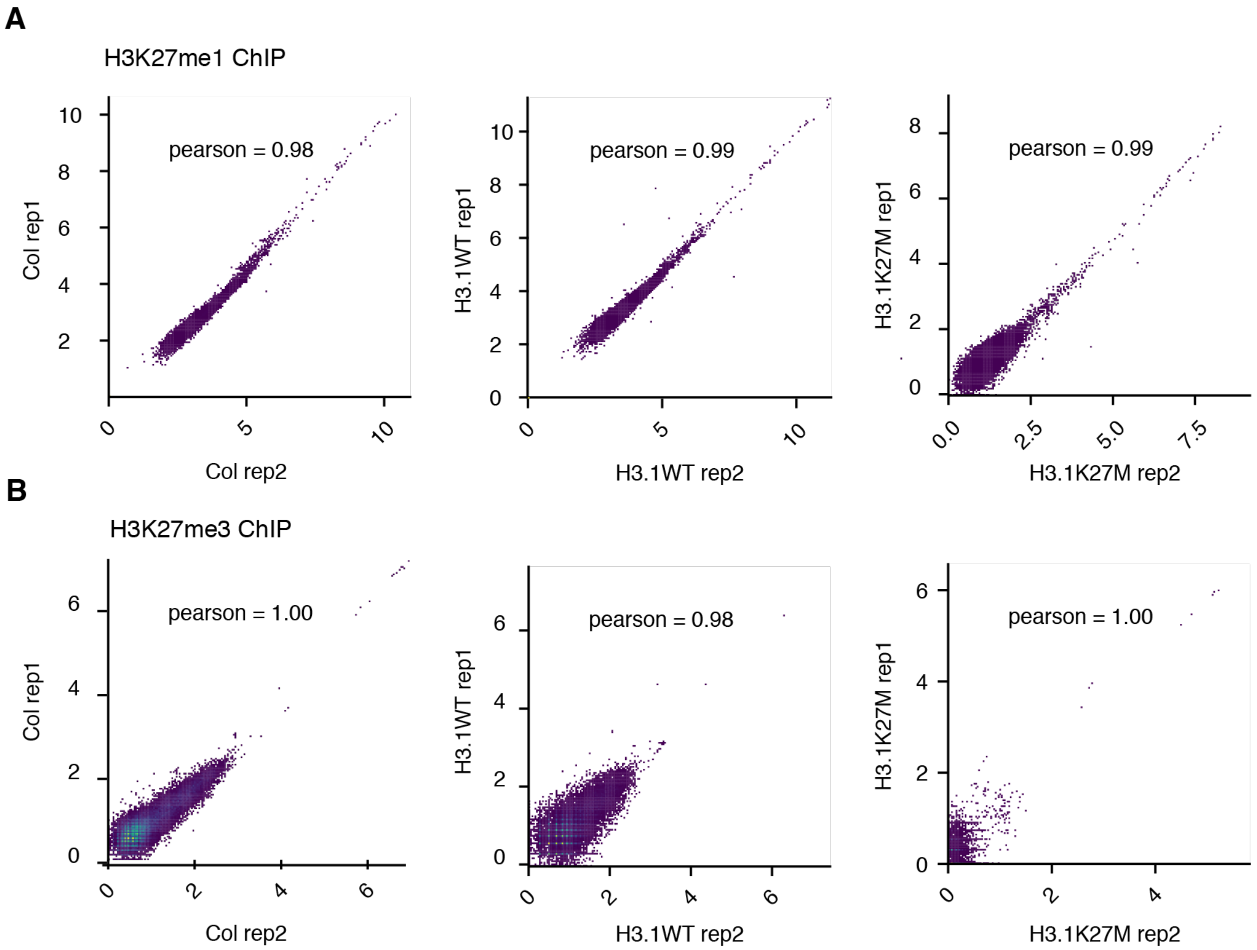


**Fig. S7. Scatterplots and Person correlation cooefficients for ChIP-seq biological replicates.**

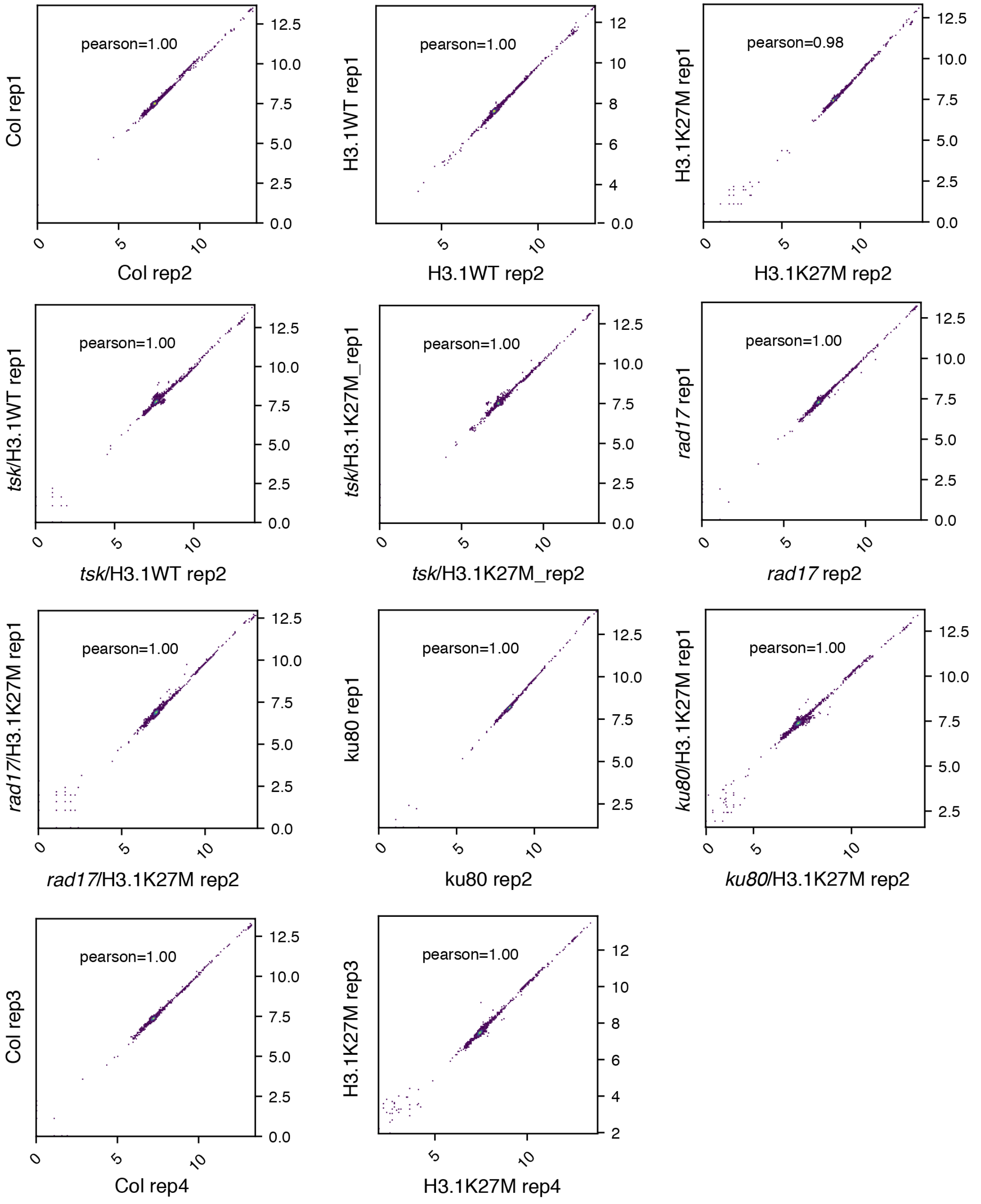


**Fig. S8. Scatterplots and Pearson correlation coefficients for DNA-seq biological replicates.**

**Table S1.**
Plant DNA repair genes.

**Table S2.**
TEs de-repressed in H3.1K27M-expressing plants.
